# Repeated evolutionary turnover of vertebrate skeletal muscle myosins

**DOI:** 10.1101/2025.10.28.684953

**Authors:** Christina M. Harvey, Eric R. Schuppe, Michael S. Brainard, Matthew J. Fuxjager, James B. Pease

## Abstract

Myosin heavy chain proteins are essential for muscle contraction and nearly every physiological function in animals, but their diversity and evolution outside mammals is largely unknown. We comprehensively model the evolutionary history of over 1100 heavy-chain myosins. We find that skeletal muscle myosins are located in a conserved tandem gene array in all vertebrate species, but repeated gene duplication-loss turnover has surprisingly led to an independently evolved set of core skeletal muscle myosins in each major vertebrate group. Despite these separate derivations of these myosin subfamilies, each major vertebrate group exhibits consistent tissue-specific patterns of subfamily expression and specialized myosin subfamily expression in extreme muscles. Our results show that muscle evolution across vertebrates is not based in conserved orthologous motor myosins, as might be expected for such a core structural protein family. Instead, we find that skeletal muscle myosins have evolved as a shifting cluster of genes that is constantly changing and diversifying to balance the need to maintain core physiology, while innovating new physiological possibilities.

## 1. Introduction

Physical movement is fundamental for animal survival and reproduction, helping individuals find food, evade predators, mate successfully, and parent offspring. Myosin heavy chain (MYH) proteins actuate much of this movement by endowing skeletal muscles with contractile properties. Physiological studies have repeatedly suggested that MYH proteins are conserved across taxa suggesting minimal evolutionary change in these core molecules [1-5]. In contrast, molecular genetic studies have found extraordinary diversity in MYH proteins broadly, and have long hinted at a more complex underlying process for the adaptive evolution of skeletal muscle MYHs involving repeated evolution of distinct sets of MYHs [6-14]. The fundamental tension between diametric perspectives of genetic conservation and repeated divergence can only be resolved by comparing MYH gene sequences and expression across a large group of unrelated taxa. Such work promises to reveal how MYH proteins have diversified historically, how they contribute to contemporary performance traits, and the central question of whether myosin evolution is dominated by slow conservative evolution or repeated divergence and innovation.

Vertebrate skeletal muscle MYH proteins pull against actin filaments to generate a power stroke that creates force by shortening or lengthening the muscle cell. MYH proteins initiate this process by hydrolyzing ATP into ADP to power actin-myosin binding kinetics [15-18]. Even subtle amino acid sequence variation can confer major alterations to MYH molecular properties [19-22]. The presence of impactful variation creates ample opportunity for selection to drive evolutionary changes in muscle performance.

Mammals have a set of sarcomeric MYH proteins known to have distinct expression patterns and molecular properties that have contributed to the past categorization of “fast” and “slow” contractile speeds [4,23-25]. MYH3 and MYH8 are primarily expressed in embryonic and neonatal tissues, with only minimal expression in some specialized adult muscles [26,27]. Adult skeletal muscles typically express MYH1, MYH2, MYH4 that are exclusively expressed in skeletal muscles, along with MYH7 that has roles in both cardiac and skeletal muscles (Fig 1A)[26]. Larger mammals tend to use a combination of MYH1, MYH2, and MYH7 in their skeletal muscles, but smaller-bodied rodents have been shown to primarily express MYH4 [28,29]. The molecular properties of these MYH proteins have been shown to be distinct with regard to ATP efficiency and mechanical movement, which connects the different MYH molecular properties to the performance properties of the muscles [21].

**Figure 1.**
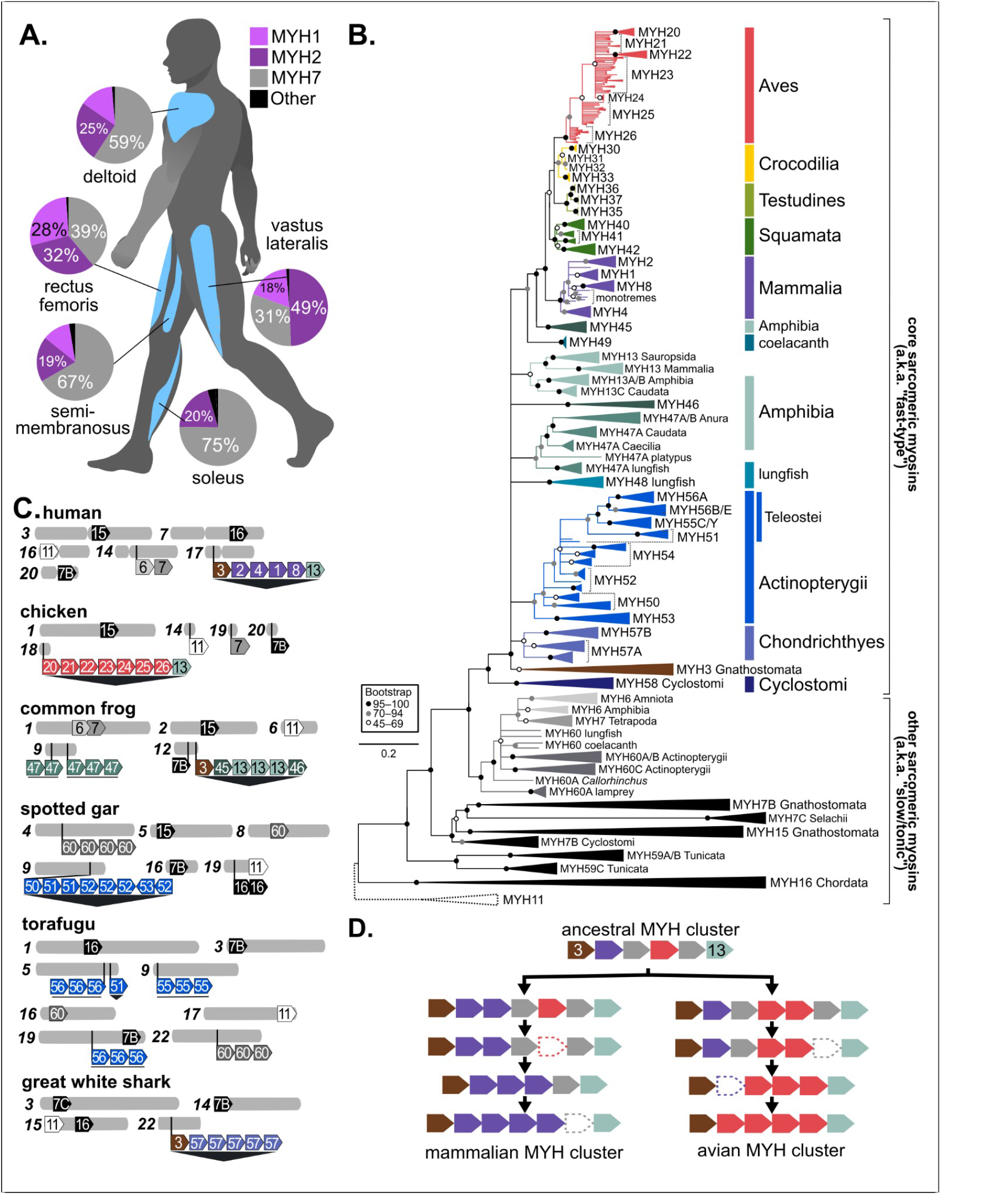

**Figure 2.**
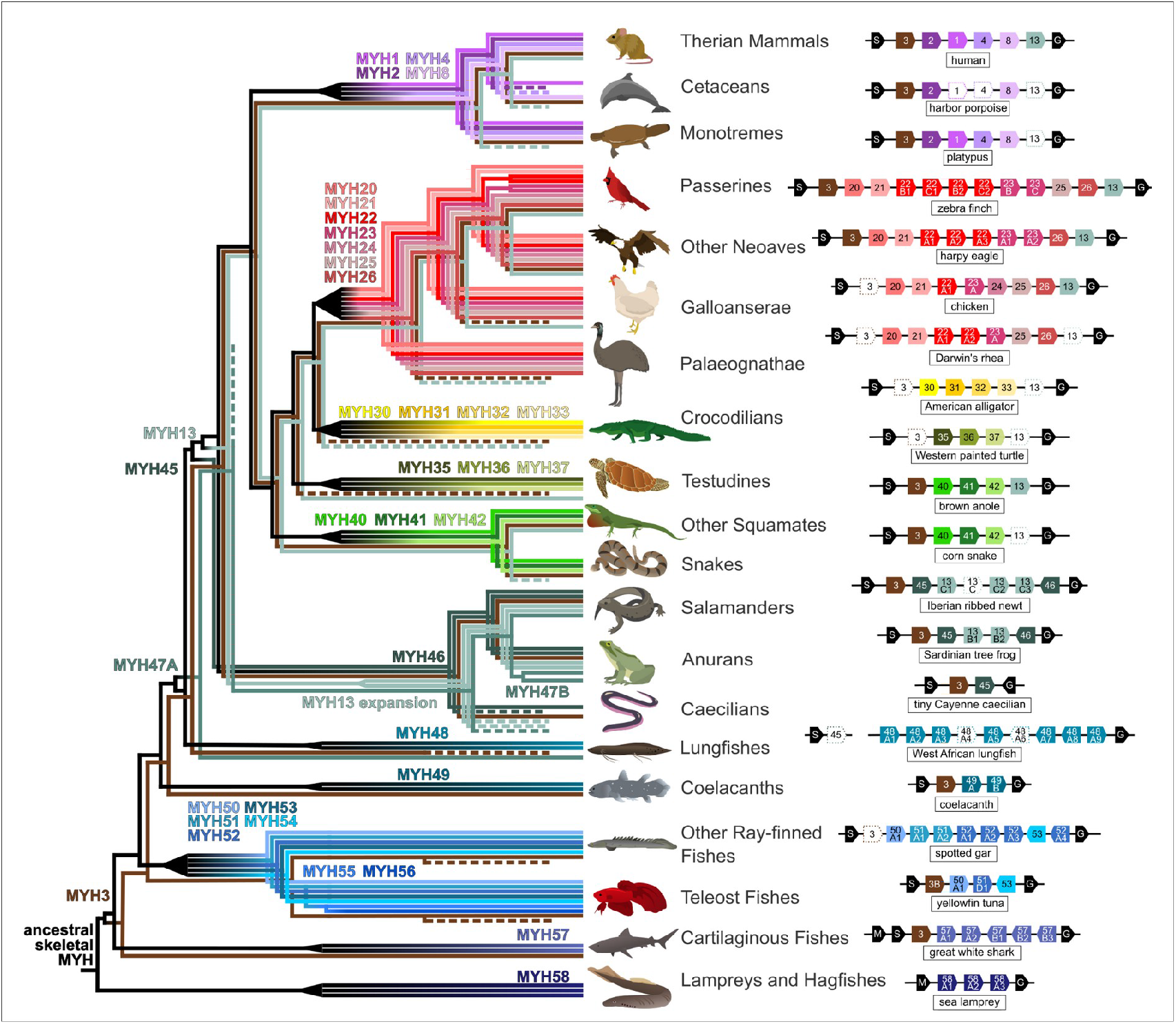
Core skeletal muscle MYH proteins are not one-to-one related among vertebrate classes. Vertebrates show repeated evidence of group-specific myosin subfamily expansion (black triangles) in their evolutionary history (left) and variation in the number of *MYH* genes in their core skeletal muscle MYH cluster (right). Specialized MYH proteins *MYH3* and *MYH13* are related across classes, but pseudogenization through mutational inactivation or partial deletion is also common (dotted outlines). Gene gain-loss in the core cluster occurs in a syntenically conserved locus near *GAS7* (G) and *SCO1* (S) or *MAP24K2* (M). The current lungfish assembly places *SCO1* next to a putatively pseudogenized *MYH45* on a separate chromosome from the rest of the cluster. (Art credit for all drawings: C. Harvey).

Current conventional wisdom holds that other vertebrate genomes contain MYHs that are orthologous in origin and function to those observed in mammalian genomes. This belief most likely stems from the observation that distinct muscle fiber types with notable performance differences have been observed in a variety of nonmammals [30,31]. For example, “superfast” muscles in vertebrates support this idea, where extremely rapid contraction-relaxation speeds are associated with distinct MYH compositions [14,32]. The presence of distinct MYHs in these tissues has been noted, though their genomic origins and relatedness to those observed in mammalian fibers have not been assessed in detail [33,34]. MYHs across vertebrates may be genetically and functionally distinct in ways that have yet to be fully described [8]Fig. 1A;. This suggests a different relationship between physiological evolution of vertebrate muscles and the molecular evolution of MYHs where evolution of individual MYH sequences and evolution of the relative expression of MYHs within muscles are joint, interrelated processes.

Here, we conduct a comprehensive study of the origins and expression of MYH genes across chordates by analyzing over a thousand MYH sequences and dozens of skeletal muscle expression profiles. We use our results to test whether MYH evolution is more generally conservative or diversifying. We find that vertebrate genomes encode sets of MYH proteins that are more diverse in gene number, genomic arrangement, and molecular structure than has been described previously. Our results conclusively demonstrate that primary skeletal MYH proteins in each major group of vertebrates evolved separately with respect to each other and lack one-to-one evolutionary relationships. These separate MYH diversifications precludes the possibility of homologous functional roles and suggests convergent pressures where similar muscle properties and functions occur in separate groups. Finally, the tissue-specific expression of MYHs in both ordinary and extreme-performance muscles points to a strong and generalized connection across vertebrates between MYH gene functional properties and muscle performance. Most broadly, this expanded evolutionary history of sarcomeric MYHs shows that skeletal myosins show lability despite being fundamental to animal reproduction and fitness, demonstrating a dramatic example of how core genes simultaneously maintain homeostasis while adapting new innovations.

## 2. Materials and methods

### (d) RNA-Seq Data

*Lonchura striata* syrinx raw RNA-seq sequences are available from NCBI SRA under BioProject #. All other expression data were obtained from public sets available on NCBI. The accession numbers to these sets are provided in Data S5 and paper DOIs are provided when available

### (e) Compiling Amino Acid Sequences

We compiled a sample of amino acid sequences by searching within the NCBI reference genomes of select species for myosins or their syntenic neighbors (Data S1 and Data S2). For species that did not have well annotated reference genomes, we sought out sequences by using NCBI’s Basic Local Alignment Search Tool for proteins (BLASTp) or expanded searches to ENSEMBL. This resulted in a set of 1397 sequences prior to filtering and quality control.

### (f) Sequence Alignment and Phylogenetic Reconstruction

We filtered our dataset to remove sequences that were partial or of poor quality. We also made small manual edits as needed to remove regions prior to start codons, retained introns, and extended repetitive ends erroneously retained on the c-terminus (see Data S4 for a description of these edits). After filtering, our dataset included 1203137 sequences from 1192 vertebrate species (Data S1 and Data S2). We aligned these sequences using MAFFT’s L-INS-i method and created phylogenetic trees using RAXML-ng with the QMaker PFAM model of protein evolution with gamma-distributed rates [36,37]. This was done with the following options:

~~~
raxml-ng --all --msa ALIGNMENT.fa --model Q.PFAM+G
~~~

RAXML-ng generates support values by default using the Felsenstein Bootstrapping protocol. Bootstrap replicates were generated until the majority rule extended (MRE) threshold was surpassed, up to a maximum of 1000 replicates.

### (g) Phylogenetic Relationships Between Myosins

We determined evolutionary relationships between myosins using both phylogenetic reconstruction and amino acid sequence analysis. First, we grouped phylogenetically monophyletic groups to identify clades of independently diverging myosins. We additionally compared sequence similarity by calculating the pairwise difference between amino acids sequences (Fig. S1). To do this, we used a custom Python3 script to count the pairwise percent distances in amino acids. The numerator was the count of all amino acid sites without a gap in both sequences that were the same amino acid. The denominator was a count of all amino acid sites without a gap in both sequences. Additionally, we computed a multidimensional scaling plot (MDS) from the same pairwise distances using the Python3 *mds* function from the *scikit-sklearn* module (Fig. S2).

We used this combined data to make the best educated hypothesis of evolutionary relatedness between myosin sequences. To differentiate between non-orthologous myosins, we renamed myosins from several animal clades to better represent their evolutionary history. These names are described in detail in Supplementary Table S1, and further detailed in the Supplementary Text.

### (h) RNA Expression Quantification and Determination of Differentially Expressed Genes

We mapped RNA expression data to NCBI reference genomes using STAR and quantified expression with featureCounts [38,39]. We began by preparing genome directories using the following options:

~~~
STAR --runMode genomeGenerate --genomeDir ./GENOMEDIR --genomeFastaFiles
GENOMEFILE.fna --sjdbGTFfile GTFFILE.gtf
~~~

We then acquired SRA data using FasterQ from the Sequence Read Archive (SRA) Toolkit and trimmed the reads by using the following command:

~~~
fasterq-dump SRA_ID --p --W --split-3
~~~

We then used STAR to align expression data against the prepared genome using:

~~~
STAR --genomeDir ./GENOMEDIR --readFilesIn SAMPLES.fq
~~~

To quantify expression data, we used featureCounts using the following command:

~~~
featureCounts --p --a GTFFILE.gtf --o OUTPUT.txt STAR_OUTPUT.bam
~~~

We transformed the raw read counts into CPM (counts per million) by dividing the counts for each gene by the total number of counts per sample and multiplying by one million. Because myosins are often very similar in sequence and sometimes differ by only a few amino acids, we removed multi-mapped reads. This means that RNA expression depicted in our data is likely lower than is actually present within the tissue, but more accurately represents the diversity and relative abundance of transcripts.

For Sika, we quantified RNA expression using *kallisto* [40]:

~~~
kallisto quant -t 64 --index $INDEX -o $OUTPUT $FASTQ1 $FASTQ2
~~~

To identify patterns of tissue specificity and changes in transcript abundance, we quantified □ and log2 maximum expression [41]. We considered genes with τ > 0.75 to exhibit high tissue-specific expression.

### (i) Calculating Conservation Score, Pairwise Distances, and Amino Acid Diversity

To determine areas of high and low conservation, we calculated conservation scores based on an alignment of 624 sequences from the tetrapod core cluster (Data S6, Data S7). Conservation scores were obtained from JalViewJS using the Analysis of Multiple Sequences (AMAS) method [51,52].

We were additionally interested in understanding instances of gene conversion or nonhomologous cross over. To identify potential regions of interest, we compared sequences we suspected of having evidence of these events to suspected orthologous sources in closely related species. We calculated pairwise distances along a sliding window (P-distance) by quantifying the proportion of nucleotide sites that differ between compared sequences (excluding gap and “N” positions when present). Distances were computed across the alignment, assigning each value to the midpoint of each window to produce a position-specific snapshot of divergence. Window sizes were assigned based on their ability to produce statistically stable P-distance estimates without compromising identification of potential recombination breakpoints.

The primary alignment (Supplementary Data S1) was filtered to four data subsets: (1) 227 sequences of MYH1–4, MYH6–8, and MYH13 from 35 mammal species, (2) 349 sequences of MYH3, MYH6, MYH7, MYH13, and MYH20–26 sequences from 36 bird species, (3) 130 sequences of MYH3, MYH6, MYH7, MYH13A, MYH13B, MYH13C, and MYH45–48 from 12 amphibian species, and (4) 106 sequences of MYH3, MYH52–55 and MYH60 from 6 ray-finned fish species. For each position in the amino acid alignment, the average Grantham’s distance for each pair of amino acids at the site was calculated as:

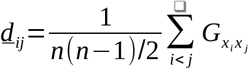

where *d*_*ij*_ is the average Gratham’s distance among all pairs of amino acids at a given position, 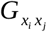 is Grantham’s distance for a given pair of amino acids from sequences *i* and *j* at position *x*, and *n* is the total number of sequences at the position [53]. Positions with more than one sequence having a gap were scored as zero. Eight sequences with partial missing sequences of >10 aa were excluded. This metric should be higher at aligned positions with both higher amino acid sequence diversity and greater amino acid side chain biochemical diversity. Calculations and plotting were performed with a custom Python3 script.

### (j) Determining Loop Lengths and Structure

Alignments of Loop 1 and Loop 2 were extracted from the primary alignment (Supplementary Data S1). We determined the start and stop of each loop based on previous work on humans and chicken myosins [3]. We calculated the ungapped length of these regions (see Data S5) and generated Logo-like plots from a sequence alignment of Loop 2 based on myosin type (Data S5, Fig 4).

## 3. Results

### (a) Core skeletal muscle MYH proteins are not one-to-one related among vertebrate classes

To create a comprehensive model of sarcomeric vertebrate MYH evolution, we modeled the molecular relationships of 1137 sarcomeric heavy-chain myosin (MYH) amino acid sequences from 113 species from all classes and most major orders of chordates (Fig. 1B). Analysis of the evolutionary change in *MYH* sequences and genomic locations reveals broader patterns of vertebrate evolution and many specific patterns within vertebrate subgroups (see Supplementary Text). We find that all vertebrate sarcomeric MYH proteins are related to a single common ancestor, with cytoskeletal *MYH11* as an outgroup. Across vertebrates, *MYH* gene copy numbers and genomic arrangements are far more variable than have been reported previously (Figs. 1B and 1C)[14].

Present across all vertebrates is a tandem cluster of 1–12 *MYH* genes, which has maintained the same syntenic context throughout vertebrate evolution). This cluster generally includes the core skeletal muscle *MYH* genes (which have historically been referred to as the “fast-type” *MYH* genes; but see Discussion) in each major group of vertebrates (Fig. 1C. While phylogenetic, pairwise distance, and MDS analyses show that embryonic MYH3 and specialized/amphibian MYH13 are related across major groups, core skeletal MYHs do not have one-to-one relationships between any two major vertebrate groups (Fig 1B; Supplementary Figs. S1 and S2)[8]. Instead, each group’s present core MYHs descend from a group-specific common ancestor (Figs. 1B, 1D, and 2) [14]. As a consequence, each major group of vertebrates has a set of *MYH* proteins that have evolved and diversified separately. Therefore, each vertebrate group’s set of MYHs would change in response to their separate evolutionary pressures. Within each group, we also identified clearly conserved and previously unidentified *MYH* subtypes. This discovery prompted us to establish a new classification for non-mammalian subtypes *MYH20–MYH60* to promote discussion of their distinct functional roles and evolutionary histories (Fig. 1B, 1C; Supplementary Tables S1 and S2).

Core skeletal cluster *MYH* genes apparently evolve via tandem gene gain and loss, where each group’s present MYH proteins are the result of gene duplications and deletions that all occurred after divergence from other groups (Figs. 1D and 2). The core skeletal cluster is a gene cassette consisting of an uninterrupted tandem array of genes that are all encoded on the same strand and by the same number of exons. We frequently observed instances of recent duplicates (>99% similar) and inactivated and partial pseudogenes. The appearance of these features in several independent lineages indicates repeated instances of gene gain and loss via unequal crossing over as the primary genetic mechanism. We further observed sequence homogenization through gene conversion, biased nucleotide content shifts, and intergenic recombination, but the evidence indicates these are lesser and rarer factors in driving cluster evolution in the long term (Supplementary Figs. S3 and S4, Supplementary Text). Amphibians, lungfish, and many ray-finned fishes have additional translocated individual MYHs or tandem clusters whose sequence relationships indicate they are derived from the core skeletal cluster (*MYH47, MYH48, MYH55* subtypes). These other tandem clusters also show expansion, contraction, and pseudogenization that indicates evolution under the same process as the core cluster. We conclude that this core skeletal MYH cluster as a whole has persisted throughout vertebrate history as a conserved gene cassette, but the specific individual genes in each cluster have gradually been replaced through duplication and loss. Additionally, translocated duplicates have established alternate clusters in certain vertebrate groups.

### (b) Vertebrates express their separate MYH protein sets in muscle-specific combinations

Expression patterns of sarcomeric MYHs have largely been studied in mammals, where correlations between MYH expression and muscle performance are commonly observed [4]. However, the findings described above that other vertebrate groups have independently evolved sets of MYHs provokes the question: do other vertebrate groups exhibit patterns of muscle-specific expression of their various MYH protein sets? Knowledge of the individual and collective patterns of expression are a window into the potential performance properties of muscles in the present and the past evolutionary regimes that have shaped them. To address this, we quantified whole RNA-Seq data sets from diverse vertebrate species where multiple muscles were available (Fig. 3). We selected cases where we could analyze locomotor muscles, specialized muscles, and extreme performance muscles. In mammals, both human and sika deer show expression of *MYH1, MYH2*, and *MYH7* in major locomotor skeletal muscles (Figs. 1A and 3A; Supplementary Fig. S5). *MYH* genes were also in the highest 1% of genes by expression level and showed moderate levels of tissue specificity in skeletal muscles. Bat breast muscles expressed *MYH1, MYH2*, and the usual cardiac-specific *MYH6*, but the “superfast” laryngeal muscles that generate echolocation signals expressed almost exclusively *MYH4* ( [54,55]. A primary locomotor muscle in the harbor porpoise expressed only *MYH2* out of the core skeletal muscle myosins, corroborating our observation that all cetaceans appear to have pseudogenized *MYH1, MYH4*, and *MYH13* (Fig. 3C).

**Figure 3.**
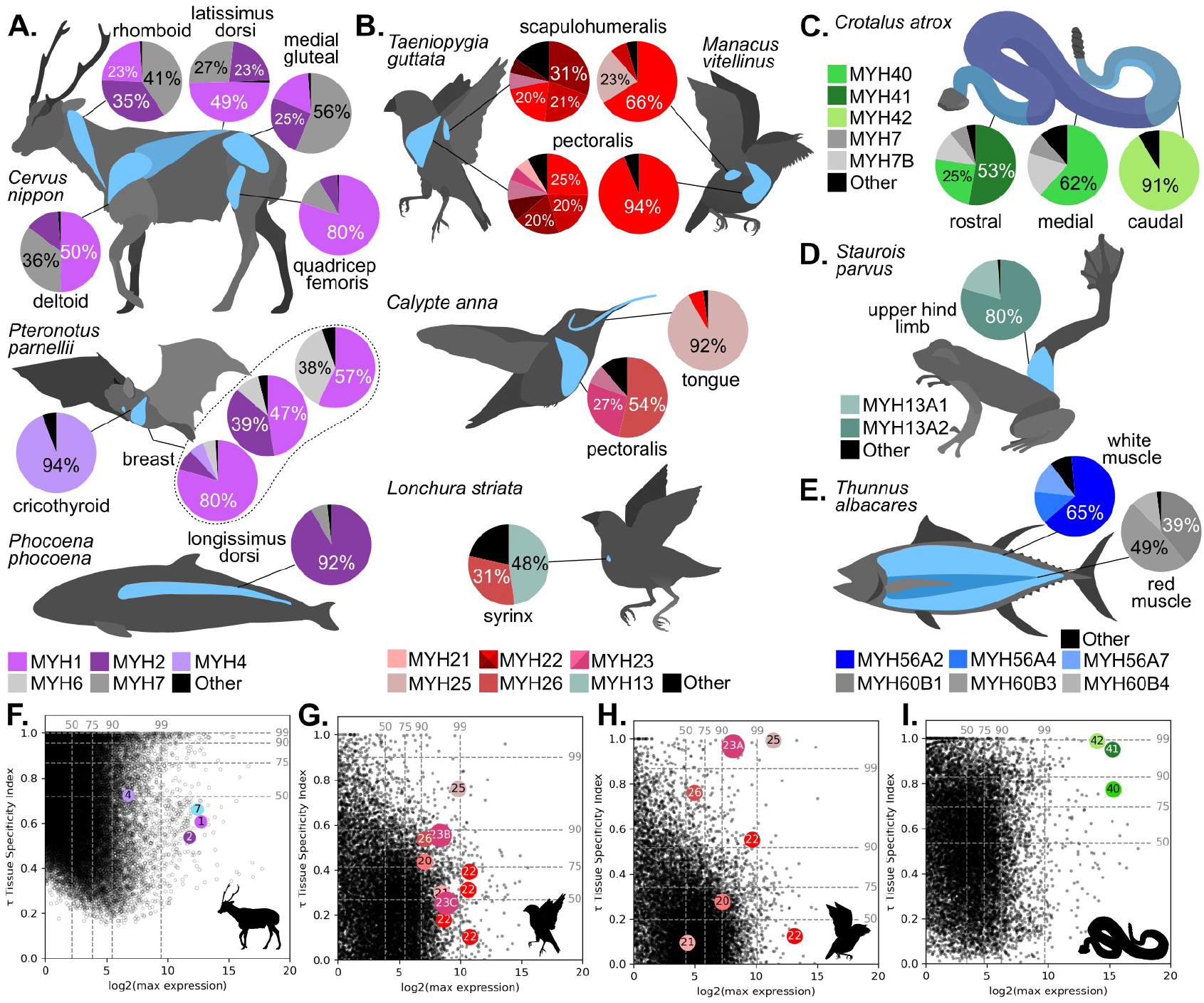
*MYH* genes show variable expression in muscles across vertebrates. The mean proportion of *MYH* expression in each of the highlighted muscles for (A) mammals (sika, Parnell’s mustached bat, and harbor porpoise), (B) birds (zebra finch, golden-collared manakin, Anna’s hummingbird, and Bengalese finch), (C) Western diamondback rattlesnake, (D) white-spotted foot-flagging frog, and (E) yellowfin tuna. *MYH* genes < 5% relative expression are included as “Other.” Breast muscles of Parnell’s mustached bat individual samples are depicted with their own pie charts. (F–I) Maximum log_2_-expression and tissue specificity index (*τ*) for all genes with *MYH* genes highlighted as color circles labeled with *MYH* numbers for (F) sika, (G) zebra finch, (H) golden-collared manakin, and (I) Western diamondback rattlesnake. Data distribution percentiles (dashed lines) are shown for each axis. (Art credit for all silhouettes: C. Harvey). See methods and supplementary information for data availability.

For birds, we focused on two passerine species, zebra finches and golden-collared manakins. Two important muscles that power flight and control wing posture, respectively, showed consistent and differentiated patterns of expression involving multiple MYHs (Fig. 3D, E). In zebra finches, the flight-powering muscle expressed seven of its twelve *MYH* genes at relative expression proportions > 5%, while golden-collared manakins overwhelmingly expressed a single *MYH22*. Notably, this latter species is known for its rapid wing-snap courtship display behavior [56]. The wing muscle that actuates this gestural signal showed expression of both *MYH22* and *MYH25*. Interestingly, this same gene (*MYH25*) is also 92% of the *MYH* expression in hummingbird flight-powering muscle (Fig. 3F). The presence of *MYH25* in two separately evolved extreme muscles strongly suggests the possibility of specialization of this *MYH25* protein. Meanwhile, “superfast” syringeal muscle from Bengalese finches showed strong expression of *MYH13*, as expected from previous evidence in zebra finches (Fig. 3G; Mead et al. 2017). However, we also found substantial syringeal expression of *MYH26*, also the most prevalent myosin in hummingbird tongues (Fig. 3F).

Other vertebrate groups show similar expression patterns. In reptiles, Western Diamondback rattlesnakes show distinct *MYH* expression patterns in three different muscle regions in its body, which govern different types of movement from locomotion to tail-shaking (Fig. 3H) [57]. Across individuals, each segment consistently showed the highest expression of a different *MYH* gene. Notably, the “superfast” caudal rattle-controlling muscles expressed high amounts of *MYH42*, which was absent from the other segments [57,58]. In amphibians, Bornean rock frogs expressed multiple forms of the *MYH13A* subtype in hind limb musculature (Fig. 3I). Amniotes have no more than one copy of *MYH13* that is used primarily in specialized muscles like the larynx and syrinx. However, amphibians have many copies of related protein subtypes MYH13A, MYH13B, and MYH13C. This case study preliminarily confirms that frogs use proteins from the MYH13 subfamily as primary locomotor myosins. Finally, an analysis of yellowfin tuna shows expression of multiple MYH55 proteins in white muscle and several MYH60 proteins from a separate cluster in red muscle (Fig 3E, Supplementary Fig. S).

Collectively, these expression data cases show that MYH proteins are not interchangeable, but instead show tissue specificity and consistent ratios among locomotor and specialized muscles. Tissue-specific MYH expression is a common theme across major vertebrate groups, which is notable given the independent evolution of their MYH gene sets. Intriguingly, this shifting balance of expression of MYH genes broadly observed across vertebrates is complementary with recent evidence that cluster MYH genes are collectively activated by a superenhancer described in mice [59]. In particular, we further show that various specialized and “superfast” muscles predominantly express different myosins, and not a universal “extreme myosin.” The tissue-specific combinations of MYH strongly suggest that these proteins have diversified to have different molecular properties, and then combine in muscles to facilitate various muscle performance properties.

### (c) Vertebrate sarcomeric myosin diversity is concentrated in two ATP-interacting loops

To explore the molecular variation of MYH proteins both within each species and across protein subfamilies, we analyzed sequence variation of core skeletal muscle MYHs, MYH3, MYH6, MYH7, and MYH13 in a structural context. MYH sequences are highly conserved within and among subfamilies across the protein, except two regions in the motor domain (Fig. 4A–C). These variable regions constitute the known loop structures “Loop 1” and “Loop 2.,” which are intermolecular binding sites for the crucial interaction with ATP. Both loops include several positively and negatively charged residues separated by multiple short amino acid segments of variable length (Fig. 4D). Across them, the overall length of the Loop 1 (11–20aa) and Loop 2 (20–31aa) vary by 2- and 1.5-fold, respectively. This length variation stands in sharp contrast to the rest of the MYH protein, where even a single amino acid insertion or deletions are virtually nonexistent except for minor 2–3aa variation in the termini. Furthermore, we found that each species has a set of MYHs with diverse Loop 1 and 2 lengths (Figs. 4E–H). Even recently duplicated subfamilies exhibit variable loop length, such as MYH22A1–MYH22A4 in pigeon (Fig. 4F). This means that, although each group’s set of core skeletal MYHs are separately duplicated from a group-specific ancestral protein, their loop lengths have repeatedly evolved variation across all species and in all vertebrate groups (Fig. 4I).

The structure of Loops 1 and 2 ATP-binding loop structural variation helps determine MYH contractile properties, largely governing rate-limiting kinetics of ATP and ADP. Previous investigations have noted sequence variation in Loops 1 and 2 across MYHs and explored how these properties may contribute to differences in sarcomere shortening and overall contractile performance binding [3,60,61]). Thus, we hypothesize that the differences we observe in MYH proteins among muscles and species contribute to variation in muscle performance. Supporting this view, we find that muscles with markedly different performance attributes express different ratios and combinations of MYH proteins. Importantly, these data are consistent with a model where the molecular structure of individual MYH genes is, at minimum, not being homogenized in the ATP-binding loops toward a single optimum. This is especially intriguing in the context of MYH proteins expressed in “superfast” muscles. Despite having a relatively similar performance of rapid contractions with little force generation, each species that maintains these muscles does so by expressing different MYH genes that share no obvious loop configuration properties. Selection for “superfast” muscle performance therefore has likely followed lineage-specific biochemical and biophysical pathways. Furthermore, muscle performance variation is not likely determined strictly by a particular molecular structure or even a particular MYH protein, but instead through the combined molecular properties and expression ratios of several MYH proteins with diverse structures.

To characterize further protein variation, we calculated amino acid diversity for each fully aligned site for core MYHs and MYH6/7/60 subfamilies separately within mammals, birds, amphibians, and ray-finned fish (Fig. 4J)[53]. Out of 1893 sites without gaps, we found that 412 (22%) were completely invariant across all sequences from these four groups with 68% of invariant sites appearing in the head domain. In contrast, 645 sites (34%) exhibited repeated amino acid variation in all four groups, while the remaining 278 (15%), 258 (14%), and 288 (15%) fully aligned sites were variable in three, two, or a single group, respectively. Sites showing variation in all four groups and variation hotspots overall were found in the N-terminal domain, the light-chain myosin binding site on the neck, and midway down the tail (1230– 1260). These intersected with several sites of known mammalian MYH7 functional impactful, while pathogenic variants were nearly always invariant across core myosins (Johnson, et al. 2021; Supplementary Table S3). No specific amino acid state was identified as being common to the myosins known to exhibit extreme muscle properties. This concurs with previous hypotheses that the specific impact of individual amino acid changes on MYH fibers or muscle performance as a whole is difficult to predict [3,62]. This analysis highlights that, particularly in the actin- and ATP-interacting myosin head region, MYH is balancing sites of extreme biochemical and structural conservation with sites that are repeatedly mutating and show variation in multiple independent groups. Therefore, structural variation is maintained in this gene family across long evolutionary distances, but the same regions have repeatedly evolved higher variation, consistent with our characterization of intermixed evolutionary pressures of homeostatic conservation and physiological innovation.

**Figure 4.**
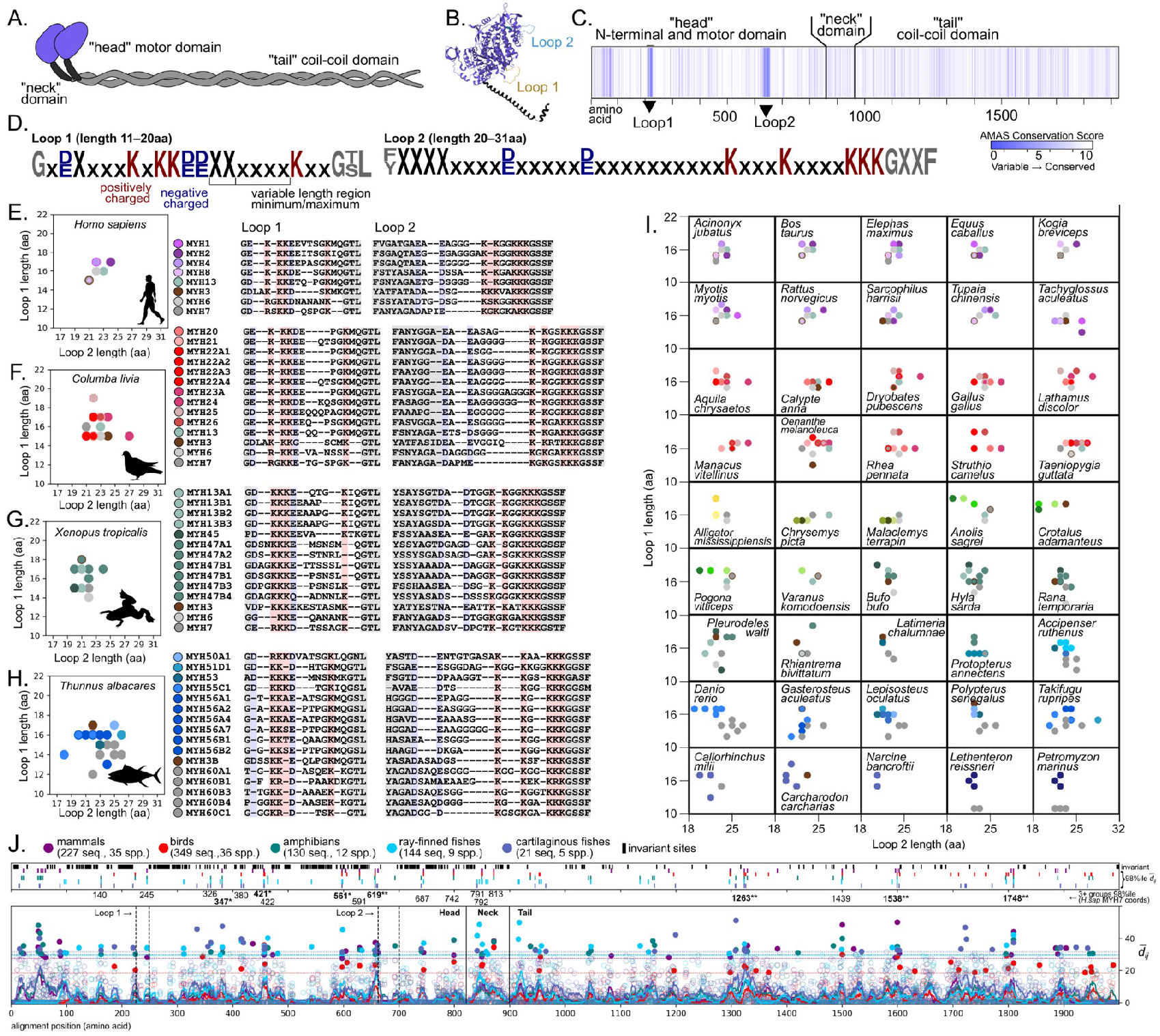
MYH loops and other sites are variable among and within vertebrate species. (A) Myosin protein structural domains shown for a dimerized protein pair. (B) Structure head and neck with Loop 1 and Loop 2 highlighted (PDB#P12882)[63,64]. (C) AMAS conservation scores for each position show that Loop 1 and Loop 2 have the highest rates of variation. (D) MYH Loop 1 and Loop 2 motifs, showing the positive and negative residues and the variable region minimum (X) and maximum (x) lengths observed. (E,F,G,H) Loop 1 and Loop 2 overall lengths create variable combinations for the core sarcomeric and slow sarcomeric MYH proteins in human, pigeon, and western clawed frog, and yellowfin tuna. (I) Loop 1 and 2 combinations for 45 additional vertebrate species show variation in MYH loop lengths is present in species in each major group. Each point represents a single MYH gene, with colors matching subfamilies in Figures 1b and 2. (J) Mean Grantham’s distance 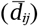 for all pairs of amino acids at each aligned position in skeletal sarcomeric myosins separately for mammals, birds, amphibians, and ray-finned fish, show amino acid positions that are recurrently variably in multiple groups. Solid graph lines show the mean value for 10aa windows. Positions with more than one gap and MYH7B, MYH15, and MYH16 were excluded. Vertical dashed lines show the Loop 1 and Loop 2 regions, and vertical solid lines show the head/heck/tail sections. Horizontal dashed lines show the 98^th^ percentiles for each group. High values 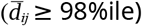 are labeled with color dots. The upper track shows the location of completely invariant sites across all sequences (black boxes) and high 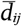 values for each group (color boxes). Numbers shown below are amino acid positions relative to human MYH7 shown for sites with high 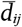 values in at least threewo groups, fourthree groups (*), or all fivefour groups (**). (Art credit for all silhouettes: C. Harvey).

## 4. Discussion

### (a) Skeletal muscle myosins evolve in a balance of functional maintenance and innovation

Understanding the molecular evolution of skeletal muscles is crucial to deciphering their physiological evolution and the evolution of vertebrates as a whole. The lack of one-to-one relationships among individual core MYH proteins between any two major vertebrate groups rejects the extrapolation of mammalian MYH functional identities or fiber type classifications to other vertebrates. emphasize an alternative perspective on skeletal muscle myosins to explain the surprising evolutionary lability of such a core genetic system. Unequal crossing over creates new myosin copies that gradually acquire diverse molecular properties (particularly in the hypervariable Loop 1 and Loop 2) leading to subfunctionalization or neofunctionalization. Specific muscles express MYH copies in specific combinations to achieve differences in performance ability, with such differences likely ranging from subtle to extreme. At the same time, many performance attributes will also require the collective action of several muscles that use a range of expression ratios of various MYH proteins. We therefore propose that MYH evolution proceeds under layers of tandem gene cluster turnover, shifting expression, and changing musculoskeletal design. This dynamic presumably creates the opportunitesy for gradual evolution, with the entire set of myosins being under a collective coselection at the phenotypic level.

### (b) An expanded view of MYH diversity prompts renewed consideration of classification

### (c) Conclusions

Several other cases of vertebrate gene family tandem cluster duplications (e.g., opsins, venom, immune factors, hormones) often involve antagonistic genetic systems whose diversification is stimulated by a shifting, exogenous factor [75-78]. The lability of MYH protein duplications and losses is therefore particularly striking, since skeletal and cardiac muscles are core, highly interconnected physiological systems presumably subject to substantial endogenous selective constraints. to understand how individual MYH genes contribute to changes in contractile force, velocity, and metabolics [79,80], this investigation establishes the rampant evolutionary turnover of myosins in the evolution of vertebrates and their independent diversifications. This new perspective on both the present diversity and underlying evolution of MYH proteins we hope will stimulate new perspectives, experiments, and discussion of the evolution of vertebrate muscles and motivate renewed investigation of the diversification of other core animal gene families.

## Ethics

Procedures involving animal use reflected in this study and not published elsewhere were conducted in accordance with institutional guidelines for the care and use of animals in research. Experimental protocols and animal care were approved by the Institutional Animal Care and Use Committee (IACUC) at the University of California of San Francisco.

## Data Accessibility

*Lonchura striata* syrinx raw RNA-seq sequences are available from NCBI SRA under BioProject #PRJNA1394504. All other data are available in the supplementary materials, or through NCBI Protein, Nucleotide, or SRA databases at the accession numbers specified therein.

## Author Contributions

Conceptualization: CMH, JBP, MJF

Methodology: CMH, ERS, JBP, MJF

Formal Analysis: CMH, JBP

Investigations: CMH, ERS, JBP

Resources: ERS, JBP, MJF, MSB

Data Curation: CMH, JBP

Writing - Original Draft: CMH, ERS, JBP, MJF

Writing - Review & Editing: CMH, JBP, MJF, MSB

Visualization: CMH, JBP

Supervision: JBP

Project Administration: JBP

Funding Acquisition: JBP, MJF, MSB

## cknowledgements

Thanks to Peri Bolton, Olivia Delgado, Lisle Gibbs, Caroline Kauh, Ellen Weinheimer, Oli Wood for feedback on this manuscript.

## References

1. Goodson HV, Warrick HM, Spudich JA. 1999 Specialized conservation of surface loops of myosin: evidence that loops are involved in determining functional characteristics. Journal of Molecular Biology 287, 173–185. (doi:10.1006/jmbi.1999.2565)

2. Weiss A, McDonough D, Wertman B, Acakpo-Satchivi L, Montgomery K, Kucherlapati R, Leinwand L, Krauter K. 1999 Organization of human and mouse skeletal myosin heavy chain gene clusters is highly conserved. Proceedings of the National Academy of Sciences 96, 2958–2963. (doi:10.1073/pnas.96.6.2958)

3. Weiss A, Schiaffino S, Leinwand LA. 1999 Comparative sequence analysis of the complete human sarcomeric myosin heavy chain family: implications for functional diversity. Journal of Molecular Biology 290, 61–75. (doi:10.1006/jmbi.1999.2865)

4. Schiaffino S, Reggiani C. 2011 Fiber types in mammalian skeletal muscles. Physiological Reviews 91, 1447–1531. (doi:10.1152/physrev.00031.2010)

5. Lee LA, Karabina A, Broadwell LJ, Leinwand LA. 2019 The ancient sarcomeric myosins found in specialized muscles. Skeletal Muscle 9. (doi:10.1186/s13395-019-0192-3)

6. Moore LA, Tidyman WE, Arrizubieta MJ, Bandman E. 1992 Gene conversions within the skeletal myosin multigene family. Journal of Molecular Biology 223, 383–387. (doi:10.1016/0022-2836(92)90741-2)

7. Moore LA, Tidyman WE, Arrizubieta MJ, Bandman E. 1993 The evolutionary relationship of avian and mammalian myosin heavy-chain genes. Journal of Molecular Evolution 36, 21–30. (doi:10.1007/bf02407303)

8. McGuigan K, Phillips PC, Postlethwait JH. 2004 Evolution of sarcomeric myosin heavy chain genes: evidence from fish. Molecular Biology and Evolution 21, 1042–1056. (doi:10.1093/molbev/msh103)

9. Foth BJ, Goedecke MC, Soldati D. 2006 New insights into myosin evolution and classification. Proceedings of the National Academy of Sciences 103, 3681–3686. (doi:10.1073/pnas.0506307103)

10. Ikeda D, Ono Y, Snell P, Edwards YJK, Elgar G, Watabe S. 2007 Divergent evolution of the myosin heavy chain gene family in fish and tetrapods: evidence from comparative genomic analysis. Physiological Genomics 32, 1–15. (doi:10.1152/physiolgenomics.00278.2006)

11. Sebé-Pedrós A, Grau-Bové X, Richards TA, Ruiz-Trillo I. 2014 Evolution and classification of myosins, a paneukaryotic whole-genome approach. Genome Biology and Evolution 6, 290–305. (doi:10.1093/gbe/evu013)

12. Wagoner JA, Dill KA. 2021 Evolution of mechanical cooperativity among myosin II motors. Proceedings of the National Academy of Sciences of the United States of America 118. (doi:10.1073/pnas.2101871118)

13. Schiaffino S, Chemello F, Reggiani C. 2024 The diversity of skeletal muscle fiber types. Cold Spring Harbor Perspectives in Biology, a041477. (doi:10.1101/cshperspect.a041477)

14. Schiaffino S, Hughes SM, Murgia M, Reggiani C. 2024 MYH13, a superfast myosin expressed in extraocular, laryngeal and syringeal muscles. The Journal of Physiology 602, 427–443. (doi:10.1113/jp285714)

15. Rayment I, Holden H, Whittaker M, Yohn C, Lorenz M, Holmes K, Milligan R. 1993 Structure of the actin-myosin complex and its implications for muscle contraction. Science 261, 58–65. (doi:10.1126/science.8316858)

16. Korn ED. 2000 Coevolution of head, neck, and tail domains of myosin heavy chains. Proceedings of the National Academy of Sciences 97, 12559–12564. (doi:10.1073/pnas.230441597)

17. Piazzesi G et al. 2007 Skeletal muscle performance determined by modulation of number of myosin motors rather than motor force or stroke size. Cell 131, 784–795. (doi:10.1016/j.cell.2007.09.045)

18. Piazzesi G, Reconditi M, Linari M, Lucii L, Sun Y-B, Narayanan T, Boesecke P, Lombardi V, Irving M. 2002 Mechanism of force generation by myosin heads in skeletal muscle. Nature 415, 659–662. (doi:10.1038/415659a)

19. Larsson L, Moss RL. 1993 Maximum velocity of shortening in relation to myosin isoform composition in single fibres from human skeletal muscles. The Journal of Physiology 472, 595–614. (doi:10.1113/jphysiol.1993.sp019964)

20. He ZH, Bottinelli R, Pellegrino MA, Ferenczi MA, Reggiani C. 2000 ATP consumption and efficiency of human single muscle fibers with different myosin isoform composition. Biophysical Journal 79, 945–961. (doi:10.1016/s0006-3495(00)76349-1)

21. Resnicow DI, Deacon JC, Warrick HM, Spudich JA, Leinwand LA. 2009 Functional diversity among a family of human skeletal muscle myosin motors. Proceedings of the National Academy of Sciences of the United States of America 107, 1053–1058. (doi:10.1073/pnas.0913527107)

22. Johnson CA et al. 2021 Identification of sequence changes in myosin II that adjust muscle contraction velocity. PLOS Biology 19, e3001248. (doi:10.1371/journal.pbio.3001248)

23. Bárány M. 1967 ATPase Activity of myosin correlated with speed of muscle shortening. The Journal of General Physiology 50, 197–218. (doi:10.1085/jgp.50.6.197)

24. Whalen RG, Sell SK, Butler-Browne G, Schwartz K, P. Bouveret, I Pinset-Härstöm. 1981 Three myosin heavy-chain isozymes appear sequentially in rat muscle development. Nature 292, 805–809. (doi:10.1038/292805a0)

25. Ennion S, Sant’ Ana Pereira J, Sargeant AJ, Young A, Goldspink G. 1995 Characterization of human skeletal muscle fibres according to the myosin heavy chains they express. Journal of Muscle Research and Cell Motility 16, 35–43. (doi:10.1007/bf00125308)

26. Schiaffino S, Rossi AC, Smerdu V, Leinwand LA, Reggiani C. 2015 Developmental myosins: expression patterns and functional significance. Skeletal Muscle 5. (doi:10.1186/s13395-015-0046-6)

27. Mascarello F, Toniolo L, Cancellara P, Reggiani C, Maccatrozzo L. 2016 Expression and identification of 10 sarcomeric MyHC isoforms in human skeletal muscles of different embryological origin. Diversity and similarity in mammalian species. Annals of Anatomy - Anatomischer Anzeiger 207, 9–20. (doi:10.1016/j.aanat.2016.02.007)

28. Lucas CA, Kang LHD, Hoh JFY. 2000 Monospecific antibodies against the three mammalian fast limb myosin heavy chains. Biochemical and Biophysical Research Communications 272, 303–308. (doi:10.1006/bbrc.2000.2768)

29. Hoh JFY. 2023 Developmental, physiologic and phylogenetic perspectives on the expression and regulation of myosin heavy chains in mammalian skeletal muscles. Journal of Comparative Physiology B-biochemical Systemic and Environmental Physiology 193, 355–382. (doi:10.1007/s00360-023-01499-0)

30. Bonine KE. 2005 Muscle fiber-type variation in lizards (Squamata) and phylogenetic reconstruction of hypothesized ancestral states. Journal of Experimental Biology 208, 4529–4547. (doi:10.1242/jeb.01903)

31. Luna VM, Daikoku E, Ono F. 2015 ‘Slow’ skeletal muscles across vertebrate species. Cell & Bioscience 5. (doi:10.1186/s13578-015-0054-6)

32. Mead AF et al. 2017 Fundamental constraints in synchronous muscle limit superfast motor control in vertebrates. eLife 6. (doi:10.7554/elife.29425)

33. Listrat A, Lebret B, Louveau I, Astruc T, Bonnet M, Lefaucheur L, Picard B, Bugeon J. 2016 How muscle structure and composition influence meat and flesh quality. The Scientific World Journal 2016, 1– 14. (doi:10.1155/2016/3182746)

34. Robinson WD, Rourke B, Stratford JA. 2021 Put some muscle behind it: understanding movement capacity of tropical birds. The Auk 138. (doi:10.1093/ornithology/ukaa068)

35. Pease JB, Driver RJ, de la Cerda DA, Day LB, Lindsay WR, Schlinger BA, Schuppe ER, Balakrishnan CN, Fuxjager MJ. 2022 Layered evolution of gene expression in ‘superfast’ muscles for courtship. Proceedings of the National Academy of Sciences 119. (doi:10.1073/pnas.2119671119)

36. Katoh K. 2005 MAFFT version 5: improvement in accuracy of multiple sequence alignment. Nucleic Acids Research 33, 511–518. (doi:10.1093/nar/gki198)

37. Kozlov AM, Darriba D, Flouri T, Morel B, Stamatakis A. 2019 RAxML-NG: a fast, scalable and user-friendly tool for maximum likelihood phylogenetic inference. Bioinformatics 35. (doi:10.1093/bioinformatics/btz305)

38. Dobin A, Davis CA, Schlesinger F, Drenkow J, Zaleski C, Jha S, Batut P, Chaisson M, Gingeras TR. 2012 STAR: ultrafast universal RNA-seq aligner. Bioinformatics 29, 15–21. (doi:10.1093/bioinformatics/bts635)

39. Liao Y, Smyth GK, Shi W. 2013 featureCounts: an efficient general purpose program for assigning sequence reads to genomic features. Bioinformatics 30, 923–930. (doi:10.1093/bioinformatics/btt656)

40. Bray NL, Pimentel H, Melsted P, Pachter L. 2016 Near-optimal probabilistic RNA-seq quantification. Nature Biotechnology 34, 525–527. (doi:10.1038/nbt.3519)

41. Yanai I et al. 2004 Genome-wide midrange transcription profiles reveal expression level relationships in human tissue specification. Bioinformatics 21, 650–659. (doi:10.1093/bioinformatics/bti042)

42. Wang Y et al. 2024 De novo transcriptome assembly database for 100 tissues from each of seven species of domestic herbivore. Scientific Data 11, 488–488. (doi:10.1038/s41597-024-03338-5)

43. Fuxjager MJ, Lee JH, Chan T, Bahn JH, Chew JG, Xiao X, Schlinger BA. 2016 Research resource: hormones, genes, and athleticism: effect of androgens on the avian muscular transcriptome. Molecular Endocrinology 30, 254–271. (doi:10.1210/me.2015-1270)

44. Borzykh AA, Makhnovskii PA, Ponomarev II, Vepkhvadze TF, Lednev EM, Rukavishnikov IV, Orlov OI, Tomilovskaya ES, Popov DV. 2025 Transcription factors associated with regulation of transcriptome in human thigh and calf muscles at baseline and after six days of disuse. Biochimica et Biophysica Acta (BBA) - Gene Regulatory Mechanisms 1868, 195086–195086. (doi:10.1016/j.bbagrm.2025.195086)

45. Osipova E et al. 2023 Loss of a gluconeogenic muscle enzyme contributed to adaptive metabolic traits in hummingbirds. Science (New York, N.Y.) 379, 185–190. (doi:10.1126/science.abn7050)

46. Ciezarek AG et al. 2018 Phylotranscriptomic insights into the diversification of endothermic Thunnus tunas. Molecular Biology and Evolution 36, 84–96. (doi:10.1093/molbev/msy198)

47. Holtz MA, Racicot R, Preininger D, Stuckert AMM, Mangiamele LA. 2023 Genome assembly of the foot-flagging frog, Staurois parvus: a resource for understanding mechanisms of behavior. G3: Genes, Genomes, Genetics 13. (doi:10.1093/g3journal/jkad193)

48. Schaiter A et al. 2024 Molecular composition of skeletal muscle in infants and adults: a comparative proteomic and transcriptomic study. Scientific Reports 14. (doi:10.1038/s41598-024-74913-4)

49. Michie KL, Kunz HE, Dasari S, Lanza IR. 2024 The influence of aging on the unfolded protein response in human skeletal muscle at rest and after acute exercise. Medicine & Science in Sports & Exercise 56, 2135–2145. (doi:10.1249/mss.0000000000003508)

50. Depuydt CE, Goosens V, Janky R, D’Hondt A, De JL, Noppe N, Derveaux S, Thal DR, Claeys KG. 2022 Unraveling the molecular basis of the dystrophic process in limb-girdle muscular dystrophy lgmd-r12 by differential gene expression profiles in diseased and healthy muscles. Cells 11, 1508–1508. (doi:10.3390/cells11091508)

51. Livingstone CD, Barton GJ. 1993 Protein sequence alignments: a strategy for the hierarchical analysis of residue conservation. Bioinformatics 9, 745–756. (doi:10.1093/bioinformatics/9.6.745)

52. Waterhouse AM, Procter JB, Martin DMA, Clamp M, Barton GJ. 2009 Jalview Version 2--a multiple sequence alignment editor and analysis workbench. Bioinformatics 25, 1189–1191. (doi:10.1093/bioinformatics/btp033)

53. Grantham R. 1974 Amino acid difference formula to help explain protein evolution. Science (New York, N.Y.) 185, 862–4. (doi:10.1126/science.185.4154.862)

54. Lee JH et al. 2018 Molecular parallelism in fast-twitch muscle proteins in echolocating mammals. Science Advances 4, eaat9660. (doi:10.1126/sciadv.aat9660)

55. Usui K, Yamamoto T, Khannoon ER, Tokita M. 2024 Musculoskeletal morphogenesis supports the convergent evolution of bat laryngeal echolocation. Proceedings of the Royal Society B Biological Sciences 291. (doi:10.1098/rspb.2023.2196)

56. Fuxjager MJ, Goller F, Dirkse A, Sanin GD, Garcia S. 2016 Select forelimb muscles have evolved superfast contractile speed to support acrobatic social displays. eLife 5. (doi:10.7554/eLife.13544)

57. Bothe MS, Kohl T, Felmy F, Gallant J, Chagnaud BP. 2024 Timing and precision of rattlesnake spinal motoneurons are determined by the KV7_2/3_ potassium channel. Current Biology 34, 286-297.e5. (doi:10.1016/j.cub.2023.11.062)

58. Martin JH, Bagby RM. 1973 Properties of rattlesnake shaker muscle. The journal of experimental zoology 185, 293–300. (doi:10.1002/jez.1401850303)

59. Dos Santos M et al. 2022 A fast Myosin super enhancer dictates muscle fiber phenotype through competitive interactions with Myosin genes. Nature Communications 13, 1039–1039. (doi:10.1038/s41467-022-28666-1)

60. Sweeney HL, Rosenfeld SS, Brown F, Faust L, Smith J, Xing J, Stein LA, Sellers JR. 1998 Kinetic tuning of myosin via a flexible loop adjacent to the nucleotide binding pocket. Journal of Biological Chemistry 273, 6262–6270. (doi:10.1074/jbc.273.11.6262)

61. Murphy CT, Spudich JA. 1999 The sequence of the myosin 50−20k loop affects myosin’s affinity for actin throughout the actin−myosin ATPase cycle and its maximum ATPase activity. Biochemistry 38, 3785–3792. (doi:10.1021/bi9826815)

62. Robert-Paganin J, Pylypenko O, Kikuti C, Sweeney HL, Houdusse A. 2019 Force generation by myosin motors: a structural perspective. Chemical Reviews 120, 5–35. (doi:10.1021/acs.chemrev.9b00264)

63. Berman H, Henrick K, Nakamura H. 2003 Announcing the worldwide Protein Data Bank. Nature Structural & Molecular Biology 10, 980–980. (doi:10.1038/nsb1203-980)

64. Sehnal D et al. 2021 Mol* Viewer: modern web app for 3D visualization and analysis of large biomolecular structures. Nucleic Acids Research 49. (doi:10.1093/nar/gkab314)

65. Bandman E, Rosser BWC. 2000 Evolutionary significance of myosin heavy chain heterogeneity in birds. Microscopy Research and Technique 50, 473–491. (doi:10.1002/1097-0029(20000915)50:6%3C473::aid-jemt5%3E3.0.co;2-r)

66. Wagner A. 1994 Evolution of gene networks by gene duplications: a mathematical model and its implications on genome organization. Proceedings of the National Academy of Sciences 91, 4387–4391. (doi:10.1073/pnas.91.10.4387)

67. Edelman GM, Gally JA. 2001 Degeneracy and complexity in biological systems. Proceedings of the National Academy of Sciences 98, 13763–13768. (doi:10.1073/pnas.231499798)

68. Schiffman JS, Ralph PL. 2021 System drift and speciation. Evolution 76, 236–251. (doi:10.1111/evo.14356)

69. Pellegrino MA, Canepari M, Rossi R, D’Antona G, Reggiani C, Bottinelli R. 2003 Orthologous myosin isoforms and scaling of shortening velocity with body size in mouse, rat, and human muscles. Journal of Physiology 546, 677–689. (doi:10.1113/jphysiol.2002.027375)

70. Taylor LD, Bandman E. 1989 Distribution of fast myosin heavy chain isoforms in thick filaments of developing chick pectoral muscle. Journal of Cell Biology 108, 533–542. (doi:10.1083/jcb.108.2.533)

71. Bandman E, Bennett T. 1988 Diversity of fast myosin heavy chain expression during development of gastrocnemius, bicep brachii, and posterior latissimus dorsi muscles in normal and dystrophic chickens. Developmental Biology 130, 220–231. (doi:10.1016/0012-1606(88)90428-9)

72. López-Unzu MA, Durán AC, Soto-Navarrete MT, Sans-Coma V, Fernández B. 2019 Differential expression of myosin heavy chain isoforms in cardiac segments of gnathostome vertebrates and its evolutionary implications. Frontiers in Zoology, 10,1–15. (doi:10.1186/s12983-019-0318-9)

73. Machida S, Masuoka R, Noda S, Hiratsuka E, Takagaki Y, Oana S, Furutani Y, Nakajima H, Takao A, Momma K. 2000 Evidence for the expression of neonatal skeletal myosin heavy chain in primary myocardium and cardiac conduction tissue in the developing chick heart. Developmental Dynamics 217, 37–49. (doi:10.1002/(SICI)1097-0177(200001)217:1<37::AID-DVDY4>3.0.CO;2-3)

74. Nord H, Burguiere A, Muck J, Nord C, Ahlgren U, von Hofsten J. 2014 Differential regulation of myosin heavy chains defines new muscle domains in zebrafish. Molecular Biology of the Cell 25, 1187–1409. (doi:10.1091/mbc.e13-08-0486)

75. Opazo JC, Hoffmann FG, Natarajan C, Witt CC, Berenbrink M, Storz JF. 2014 Gene turnover in the avian globin gene families and evolutionary changes in hemoglobin isoform expression. Molecular Biology and Evolution 32, 871–887. (doi:10.1093/molbev/msu341)

76. Ramirez M, Pairett A, Pankey M, Serb J, Speiser D, Swafford A, Oakley T. 2016 The last common ancestor of most bilaterian animals possessed at least 9 opsins. Genome Biology and Evolution 8, evw248. (doi:10.1093/gbe/evw248)

77. Storz JF. 2016 Gene duplication and evolutionary innovations in hemoglobin-oxygen transport. Physiology 31, 223–232. (doi:10.1152/physiol.00060.2015)

78. Mason AJ, Holding ML, Rautsaw RM, Rokyta DR, Parkinson CL, Gibbs HL. 2022 Venom gene sequence diversity and expression jointly shape diet adaptation in pitvipers. Molecular Biology and Evolution 39. (doi:10.1093/molbev/msac082)

79. Seaborne RA, Ochala J. 2023 The dawn of the functional genomics era in muscle physiology. The Journal of Physiology 601, 1343–1352. (doi:10.1113/jp284206)

80. Martínez Mir C, Pisterzi P, De Poorter I, Rilou M, van Kranenburg M, Heijs B, Alemany A, Sage F, Geijsen N. 2024 Spatial multi-omics in whole skeletal muscle reveals complex tissue architecture. Communications Biology 7. (doi:10.1038/s42003-024-06949-1)

