## Supplementary Text, Figures, and Tables for "Repeated evolutionary turnover of vertebrate skeletal muscle myosins"

**The PDF files includes:**

Supporting Text
Figs. S1 to S6
Tables S1 to S5
Supplementary References

**Other supplementary materials for this manuscript include the following:**

Data S1-S8

**Supplementary Table S1. Myosin labels and cluster/gene locations**

| Locus | Genes/Subfamilies | Clade[†] | Alternative Names[‡] |
|---|---|---|---|
| **Core Skeletal Sarcomeric Myosins** | | | |
| _GAS7_ cluster | _MYH1, MYH2_, _MYH4_, _MYH8_ | Mammalia | 2X, 2A, 2B, Neonatal |
| | _MYH20_*, _MYH21_, _MYH22_*, _MYH23_*, _MYH24_, _MYH25_*, _MYH26_ | Aves | – |
| | _MYH30_, _MYH31_, _MYH32_, _MYH33_ | Crocodilia | – |
| | _MYH35_, _MYH36_, _MYH37_ | Testudines | – |
| | _MYH40_*, _MYH41_*, _MYH42_ | Squamata | – |
| | _MYH13A_*,_MYH13B_* | Anura | |
| | _MYH13C_* | Caudata | |
| | _MYH45_ | Amphibia | – |
| | _MYH46_ | Anura, Caudata | – |
| | _MYH48_* | lungfish | – |
| | _MYH49_* | coelacanth | – |
| | _MYH50_*, _MYH51*, MYH52*, MYH53_, _MYH54A_* | Actinopterygii | – |
| | _MYH57_* | Chondrichthyes | – |
| | _MYH58_* | Cyclostomi | – |
| | _MYH3_ | Gnathostomata | Embryonic |
| | _MYH13_ | Tetrapoda | Extraocular |
| _TBC1_ cluster | _MYH47A_* | Anura, Caudata, lungfish (pseudogene in platypus) | – |
| _TKTL_ cluster | _MYH47B_* | Anura | – |
| _XAB2_ cluster | _MYH56A_* | Teleostei | "Cluster A" |
| _P4HTMB_ cluster | _MYH56B_* | Teleostei | "Cluster B" |
| _HYDIN_ cluster | _MYH55C_* | Teleostei | "Cluster C" |
| _GAS7A cluster_ | _MYH51D_* | Teleostei | "Cluster D" |
| Other fish clusters | _MYH56E*, MYH54B/C, MYH55Y_* | Actinopterygii | – |
| **Other Sarcomeric Myosins** | | | |
| _NDGN_ cluster | _MYH6_* | Tetrapoda | α-Cardiac |
| | _MYH7_* | Tetrapoda | β-Cardiac |
| | _MYH60A_* | Non-Tetrapod Vertebrates | – |
| _PRF1/MAP4L_ cluster | _MYH60B_* | Actinopterygii | – |
| _PAK6B_ retrogene | _MYH60C1_ | Actinopterygii | – |
| _CIP2A_ | _MYH15_ | Gnathostomata | Slow B |
| _TRPC4AP-GSS_ | _MYH7B_* | Vertebrata | Slow A |
| _SAMHD1_ cluster, _SENP1_ cluster | _MYH59A/BC_ | Tunicata | – |
| _KPNA7_ | _MYH16_ | Chordata | Masticatory |
| | _MYH11_ | Chordata | Smooth Muscle |

*These myosin subtypes can have more than one paralog represented in a genome in some lineages.
†Taxonomic scope to be the best of our present knowledge, but not necessarily a definitive declaration of origin.
‡Labels used in past literature that are largely based on primate, rodent, chicken, zebrafish, and takifugu models.

#### Supplementary Table S2. Additional synonyms for MYH labels

| | | |
|---|---|---|
| *MYH1* | Mammalia | Myosin Heavy Chain IIx, Myosin Heavy Chain D, MyHC-2X, MyHC-2X/D, MYH-IIx, MYHa, MYHSA1, HEL71 |
| *MYH2* | Mammalia | IBM3, CMYO6, CMYP6, MYH2A, MYPOP, MYHSA2, MYHas8, MyHC-2A, MyHC-IIa |
| *MYH3* | Mammalia | DA8, DA2A, DA2B, DA2B3, HEMHC, SMHCE, MYHSE1, CPSFS1A, CPSFS1B, CPSKF1A, CPSKF1B, MYHC-EMB, embryonic myosin heavy chain |
| *MYH4* | Mammalia | MYH2B, MyHC-2B, MyHC-IIb |
| *MYH6* | Tetrapoda | ASD3, CMD1EE, CMH14, MYHC, MYHCA, SSS3, alpha-MHC, myosin, heavy chain 6, cardiac muscle, alpha, myosin heavy chain 6, MHC-α, slow α-cardiac |
| *MYH7* | Tetrapoda | CMD1S, CMH1, MPD1, MYHCB, SPMD, SPMM, myosin, heavy chain 7, cardiac muscle, beta, myosin heavy chain 7, MHC-β, slow type I MHC, slow β-cardiac |
| *MYH7B* | Vertebrata | MHC14, MYH14, lncMYH7b |
| *MYH7B1* | Actinopterygii | myh7ba (*D. rerio)* |
| *MYH7B2* | Actinopterygii | Myh7bb (*D. rerio)* |
| *MYH8* | Mammalia | DA7, MyHC-pn, gtMHC-F, MyHC-peri, perinatal myosin heavy chain, neonatal myosin heavy chain |
| *MYH13* | Tetrapoda | MyHC-eo, MyHC-IIL, extraocular myosin heavy chain, laryngeal myosin II |
| *MYH16* | Vertebrata | MYH16P, MHC20, MYH5 |
| *MYH20* | Aves | MYH1G (*G. gallus*) |
| *MYH21* | Aves | MYH1C (*G. gallus*) |
| *MYH22* | Aves | MYH1E (*G. gallus*) |
| *MYH23* | Aves | MYH1B (*G. gallus*) |
| *MYH24* | Aves | MYH1A (*G. gallus*) |
| *MYH25* | Aves | MYH1F (*G. gallus*) |
| *MYH26* | Aves | MYH1D (*G. gallus*), chicken embryonic |
| *MYH55B1* | Actinopterygii | myhb (*D. rerio)* |
| *MYH55C1* | Actinopterygii | myhz2 (*D. rerio)* |
| *MYH55C2* | Actinopterygii | myhc4 (*D. rerio)* |
| *MYH55C3* | Actinopterygii | myha (*D. rerio)* |
| *MYH55C4* | Actinopterygii | myhz1.3 (*D. rerio)* |

| | | |
|---|---|---|
| *MYH55C5* | Actinopterygii | myhz1.2 (*D. rerio)* |
| *MYH55C6* | Actinopterygii | myhz1.1 (*D. rerio)* |
| *MYH55E2* | Actinopterygii | myhz1.1l (*D. rerio)* |
| *MYH60A3* | Actinopterygii | smyhc3 (*D. rerio)* |
| *MYH60A4* | Actinopterygii | smyhc2 (*D. rerio)* |
| *MYH60A5* | Actinopterygii | smyhc1 (*D. rerio)* |
| *MYH60B1* | Actinopterygii | myh7l (*D. rerio)* |
| *MYH60B2* | Actinopterygii | myh7 (*D. rerio)* |
| *MYH60C1* | Actinopterygii | myh6 (*D. rerio)* |

**Supplementary Table S3. Diversity at known functional/pathogenic variation sites**

| Protein Variant | Type | Diversity in MYH alignment | Reference |
|---|---|---|---|
| p.T178I (*Hs* MYH2) | human pathogenic | Invariant in all MYH | [1] |
| p.A236T (*Hs* MYH2) | human pathogenic | Invariant in all MYH | [1] |
| p.F287V (*Hs* MYH3) | human pathogenic | Y in some MYH7 and most MYH7B | [2] |
| p.E321G (*Ec* MYH1) | equine pathogenic | V in MYH7B, MYH15, MYH16<br>Q in Passeriformes MYH3<br>Q in MYH55C,F,G and some MYH52,53 | [3] |
| p.A326P (*Hs* MYH7) | human pathogenic | Highly variable, MYH-type specific | [4] |
| p.L342Q (*Mm* MYH4) | murine pathogenic | Invariant in all MYH | [5] |
| p.S343P (*Hs* MYH7) | mammal variant | Highly variable, MYH-type specific | [4] |
| p.M349I (*Hs* MYH7) | mammal variant | Mostly I in core MYH, V in other | [4] |
| p.L366Q (*Hs* MYH7) | mammal variant | Q in all except some mammal MYH7 | [4] |
| p.I421A (*Hs* MYH7) | mammal variant | Highly variable, MYH-type specific | [4] |
| p.T424I (*Hs* MYH7) | mammal variant | R or K MYH7 in artiodactyl/elephant<br>V or I in all other MYH types | [4] |
| p.A430S (*Hs* MYH7) | mammal variant | S or A variable across types<br>G in Anura MYH15 | [4] |
| p.R434K (*Hs* MYH7) | mammal variant | R in MYH7B,MYH15. K in all others. | [4] |
| p.F553Y (*Hs* MYH7) | mammal variant | F in mammal MYH15 and some avian MYH6. All others Y. | [4] |
| p.V561I (*Hs* MYH7) | mammal variant | Highly variable, MYH-type specific | [4] |
| p.P573Q (*Hs* MYH7) | mammal variant | Highly variable, MYH-type specific | [4] |
| p.I580V (*Hs* MYH7) | mammal variant | G/Q in MYH56,MYH57. All others I/V | [4] |
| p.V609M (*Hs* MYH1) | human pathogenic | L in bats and some MYH55<br>I in MYH20–24, MYH30–33<br>T in some MYH56 | [6] |
| p.R674Q (*Hs* MYH8) | human pathogenic | Invariant in all MYH | [7] |
| p.A850T (*Hs* MYH7) | human pathogenic | Q in some MYH13A/B,MYH3, others<br>S in some MYH7B | [8] |
| p.M852R (*Hs* MYH7) | human pathogenic | Cognate I/L variants common<br>K in MYH16 | [8] |
| p.K865R (*Hs* MYH7) | human pathogenic | A/T in MYH7B, some MYH55<br>D/E in Gnathostome MYH16<br>R in most mammal MYH13 | [8] |
| p.S866P (*Hs* MYH7) | human pathogenic | A in mammal MYH7B<br>A in most MYH55<br>T in great ape MYH4 and MYH1<br>A in other mammal MYH1 | [8] |
| p.V970I (*Hs* MYH2) | human pathogenic | Invariant in all MYH | [9] |
| p.L1061V (*Hs* MYH2) | human pathogenic | Invariant in all MYH | [9] |

| | | | |
|---|---|---|---|
| p.R1500P (*Hs* MYH7) | human pathogenic | Invariant in all MYH | [10] |
| p.K1617del (*Hs* MYH7) | human pathogenic | Invariant in all MYH | [11] |
| p.L1706P (*Hs* MYH7) | human pathogenic | Invariant in all MYH | [10] |
| p.K1729del (*Hs* MYH7) | human pathogenic | Invariant in all MYH | [12] |
| p.L1793P (*Hs* MYH7) | human pathogenic | Invariant in all MYH | [13] |
| p.R1845W (*Hs* MYH7) | human pathogenic | Invariant in all MYH | [13] |
| p.H1901L (*Hs* MYH7) | human pathogenic | Invariant except two rare R alleles | [13] |

**Supplementary Table S4: Genome assemblies and mapping sources for RNA expression data**

| Species Sampled | Mapped Species | Genome Assembly Accession Number | Source of Mapping |
|---|---|---|---|
| *Cervus nippon* | *Cervus canadensis* | GCA_019320065.1 | this study |
| *Staurois parvus* | *Staurois parvus* | GCA_951230385.1 | this study |
| *Homo sapiens* | *Homo sapiens* | GCA_000001405.29 | GSE257558, GSE202745, GSE271607, GSE249921 |
| *Calypte anna* | *Calypte anna* | GCA_003957555.2 | this study |
| *Taeniopygia guttata* | *Taeniopygia guttata* | GCA_003957565.4 | this study |
| *Manacus vitellinus* | *Manacus vitellinus* | GCA_001715985.3 | this study |
| *Crotalus atrox* | *Crotalus adamanteus* | GCA_039797435.1 | this study |
| *Lonchura striata* | *Lonchura striata* | GCA_005870125.1 | this study |
| *Pteronotus parnellii* | *Pteronotus mesoamericanus* | GCA_021234165.1 | this study |
| *Phocena phocena* | *Phocena phocena* | GCA_963924675.1 | this study |
| *Thunnus albacares* | *Thunnus albacares* | GCA_914725855.1 | this study |

**Supplementary Table S5: RNA-Seq quantification metrics for *Lonchura striata* syrinx reads**

| Sample | SRA | Total Reads | UniqueMap | %UniqueMap | Assigned | %Assigned |
|---|---|---|---|---|---|---|
| **bu52bu23** | SRR36609005 | 176,549,332 | 146,993,567 | 83.3% | 104,324,602 | 71.0% |
| **gr15bu53** | SRR36609004 | 225,893,536 | 203,565,574 | 90.1% | 179,293,227 | 88.1% |
| **gr34bu97** | SRR36608997 | 139,701,960 | 121,375,936 | 86.9% | 107,194,477 | 88.3% |
| **or100pu98** | SRR36608996 | 139,265,165 | 124,019,456 | 89.1% | 108,569,559 | 87.5% |
| **or13or14** | SRR36608994 | 201,993,461 | 179,482,116 | 88.9% | 158,420,231 | 88.3% |
| **or93rd47** | SRR36608992 | 201,372,022 | 179,929,784 | 89.4% | 158,696,853 | 88.2% |
| **or15or21** | SRR36608993 | 117,461,797 | 105,503,297 | 89.8% | 93,716,729 | 88.8% |
| **or11pu23** | SRR36608995 | 118,085,269 | 103,697,668 | 87.8% | 91,062,394 | 87.8% |
| **pk25rd67** | SRR36608990 | 133,718,887 | 119,087,907 | 89.1% | 105,279,115 | 88.4% |
| **pk80pk39** | SRR36608998 | 172,000,372 | 154,196,125 | 89.6% | 135,391,638 | 87.8% |
| **pk48pu45** | SRR36609003 | 160,458,072 | 141,624,643 | 88.3% | 124,152,924 | 87.7% |
| **pk21rd11** | SRR36608991 | 267,750,074 | 237,383,172 | 88.7% | 209,432,449 | 88.2% |
| **pu30pu91** | SRR36609002 | 251,452,996 | 219,242,424 | 87.2% | 193,289,030 | 88.2% |
| **rd86gr89** | SRR36609000 | 171,422,051 | 155,251,330 | 90.6% | 136,575,775 | 88.0% |
| **rd9gr38** | SRR36608999 | 224,097,091 | 200,745,174 | 89.6% | 177,081,407 | 88.2% |
| **rd48rd89** | SRR36609001 | 120,620,602 | 104,811,161 | 86.9% | 93,176,887 | 88.9% |

#### Supplementary Text

##### 1) Additional details on general patterns of gene and genomic configuration

###### 1.1) Gene Structure

Unless specifically noted below, *MYH* genes have 39–41 exons. The coding sequence (CDS) is contained nearly universally on 39 exons, and then some *MYH* genes have an additional exon that only includes 5′- or 3′-UTR sequence (no codons). The lengths of the exons are almost uniformly conserved across all taxa and forms of myosin, with the notable exception of the two variable loops (Loop 1 and Loop 2) and the 5′- or 3′-UTRs.

###### 1.2) Tandem Cluster Structure

*MYH* tandem clusters usually consist of genes on the same strand, with the exception of some of the teleost fish clusters that can sometimes have genes on both strands. This is consistent with the generally more diverse number and structure of *MYH* genes and clusters in teleosts, likely due to the whole-genome duplication and reshuffling of chromosomes in this group.

We observe that older, single-copy *MYH* genes in the cluster tended to occur at the ends of clusters, particularly the 3′ end. Examples include: *MYH3/MYH13* in amniotes, *MYH45/MYH46* in amphibians, and *MYH53/MYH54* in ray-finned fish. In contrast, more recent duplicates seem to occur in the middle of the cluster. For example, *MYH22* and *MYH23* subfamilies in birds, *MYH13B/C* in amphibians, and *MYH52* in non-teleost ray-finned fish. We propose this might be consistent with unequal crossing over as a primary mechanism, as recombination slippage would be more likely in the middle of a repeat rather than at the periphery. However, the genes at the periphery also seem to be more specialized in function, so there may also be a functional reason for their maintenance.

###### 1.3) Potential correlation between number of MYH genes and complexity of musculature

We present a general hypothesis that individuals are using different MYH proteins in different expression ratios in each muscle to achieve various performance properties. A logical extension arising from this is a hypothesis that having more numerous MYH proteins would be a necessary prerequisite to having more diverse MYHs and therefore greater potential muscle functional specificity. We only observe a single core functional MYH in caecilians and cetaceans, with the presence of several pseudogenes in both cases indicating this paucity is the result of MYH paralog attrition relative to their ancestors. Both lineages have also experienced substantial body plan changes relative to their ancestor, particularly loss of limbs and shift to aquatic habitats. However, snakes have a fairly typical number of MYH for squamates and pinnipeds and manatee have the standard mammal number and configuration. Therefore, neither a shift to a marine habitat nor loss of limbs has direct predictive power on its own. We should also be careful not to underestimate the substantial muscular performance demands on caecilian and cetacean physiology. At the other extreme, birds, frogs, salamanders, and teleost fish tend to have a large number. The demands of flight and endothermy, jumping, or fast swimming in a marine habitat could require additional muscle specialization, but we acknowledge that these correlations are vague and the variety of expression profiles possible with even a few MYHs can be profound. Therefore, we note this hypothesis for future investigation, though the evidence at this point is not sufficient to make a broad-scale determination. Since each group's myosin set is evolving independently, we will likely have to do more targeted sampling within groups for broader patterns rather than between groups.

#### 2) Additional Details on MYH Genes Orthologous Across Groups

##### 2.1) MYH16

*MYH16* is orthologous across all vertebrates, and syntenically linked to the *KPNA7* gene throughout. This gene is deleted or apparently pseudogenized in many lineages scattered throughout vertebrates. Historically, it has been known as the "mandibular myosin" since it has this role in many mammals (though notably not humans). But its presence in jawless vertebrates and tunicates and frequent losses in jawed vertebrate lineages point to a different origin and more diverse role for MYH16 across vertebrates. Based on phylogenetic evidence, it is mostly likely that *MYH16* may represent the original paralog subfamily among all other myosins that share a common ancestor with it. Two additional copies are present in *Myxine* as *MYH16B* and *MYH16C*.

##### 2.2) MYH7B

*MYH7B* is orthologous across vertebrates and generally stable in its syntenic gene neighborhood that we label the *TRPC4AP*-linked locus. An additional paralog called *MYH7B2* appear in teleost fish associated with a whole-genome duplication and a separate duplicate in *Myxine*. An additional paralog specific to Selachii (sharks) is observed and labeled *MYH7C*.

##### 2.3) MYH15

*MYH15* appears to be Gnathostome-specific and is orthologous across classes and also appears in a syntenic conserved neighborhood that we label the *CIP2A*-linked locus. *MYH15* appears to be absent in many lineages. For example, *MYH15* is present in non-teleost bony fish, but absent in all teleosts. *MYH15* is present in the lone available Holocephali genome (*Callorhincus millii*), but absent in all Elasmobranchii.

##### 2.4) The NDGN-linked locus and derivatives (MYH6/MYH7/MYH60 subfamily)

*MYH6* and *MYH7* exist as a tandem pair of single-copy orthologs in mammals at a conserved syntenic locus that we label the *NDGN*-linked locus. Outside of mammals, they can be higher in copy number, translocated, and unlinked. In birds specifically, we note that *MYH6* and *MYH7*-like genes frequently appear in various copy numbers and at unique locations within the genome. Paleognathae (ratites) maintain *MYH6* and *MYH7* near *NDGN*. In contrast, most Gallanseriform and Neoaves genome assemblies currently show either *MYH6* or *MYH7* present and the position of these loci are highly variable relative to syntenic blocks. Passerines nearly all possess a pair of *MYH6* and *MYH7* at the same locus near *UPP1* and *MED10*, though they possess an additional gene (*MYH6B*) at another location within the genome. Patterns in other avian groups appear highly variable.

We additionally note that *MYH6* and *MYH7* clades appear to only be orthologous across tetrapods. Therefore, we adopt *MYH60* to refer to ancestrally related to *MYH6* and *MYH7* that duplicated prior to the divergence of tetrapods. We further refer to these genes as *MYH60A*, *MYH60B*, and *MYH60C* to represent distinct evolutionary lineages in nontetrapods. We use *MYH60A* to refer to genes appearing at a syntenic locus as *MYH6* and *MYH7* in mammals. In some teleosts (i.e., *Thunnus*) this cluster still remains proximal to *NDGN*. In others, the genes are proximal to *CMTM5* (*Protopterus*), another syntenic neighbor observed beside the conserved *MYH6* and *MYH7* loci in mammals. In other fishes, genomic arrangements vary, though these genes still exhibit phylogenetic support as part of the *MYH60A* clade observed in syntenically conserved genomes. *MYH60B* was adopted for derived genes in this subfamily appearing at loci linked to *PRF1* (*Acipenser*) and *MAP4L* (teleosts). *MYH60C1* was found in spotted gar, torafugu, and zebrafish as a single-exon putatively retrotransposed gene linked to *PAK6B* in all three species. This was the only case of retrotransposition of an MYH observed.

Among vertebrate genome assemblies surveyed, only hagfish (*Myxine glutinosa*) had no sequenced genes from the 6/7/60 subfamily, while lampreys have several *MYH60* copies.

##### 2.5) MYH3

*MYH3* appears as part of the core skeletal *GAS7*-linked cluster and is orthologous across jawed vertebrates, but missing in many species and whole groups (especially in Sauropsida, see details below).

##### 2.6) MYH13

*MYH13* appears to be tetrapod-specific. *MYH13* in amniotes is a single orthologous locus. *MYH13* exists in higher copy number in amphibians (see details below). None of the amphibian *MYH13A, MYH13B,* and *MYH13C* genes shares a notably closer sequence distance or relationship to amniote *MYH13.*

#### 3) Details on group- and species-specific *MYH* gene sets and genomic configurations and the MYH nomenclature

##### 3.1) Mammalia

Mammals have a core *GAS7*-linked tandem cluster of *MYH3-MYH2-MYH1-MYH4-MYH8-MYH13* [14]. This configuration is constant across all therian and metatherian mammalian families, except monotremes and cetaceans. All cetaceans appear to have pseudogenized/nonfunctional *MYH1, MYH4, MYH8,* and *MYH13* sequences. This leaves only *MYH2* as their primary adult skeletal muscle myosin. Monotremes (platypus and echidna) both have a partial *MYH13* psesudogene sequence, and platypus has a visible pseudogene of *MYH47* next to the *TKTL2* gene that is orthologous and syntenic with the *MYH47* cluster in Amphibia and Dipnoi. The functional status of *MYH8* in platypus is not known, and this may also be pseudogenized. Mammals all have a tandem *MYH6* and *MYH7* locus and other *MYH* genes at the usual loci.

Mammals are notable among vertebrate groups for having such a consistent set of myosins that appear to be long-standing mammalian orthologs that are more diverged in sequence from each other compared to the tandem genes in other groups. We examined mammal genome assemblies beyond our dataset across all families and major subfamilies and found no additional duplicates of *MYH* genes in any group. Speculatively, if any amniote group has retained a configuration similar to the amniote ancestor, it would most likely be mammals. All Sauropsida groups show evidence of much more recent duplication/replacement of their *MYH* genes. However, it seems probably more likely that the mammalian set is equally derived from a class-specific ancestor gene as the rest of the vertebrate groups.

##### 3.2) Aves

Aves shows substantial paralog number variation in their *GAS7*-linked core cluster, though the number of paralogs has noticeable modes within some major clades (Figure 1)[15]. Ratites (Palaeognathae) tend to have 6–7 core *MYH20–MYH26* genes and have pseudogenized or absent *MYH3* and *MYH13*. Galloanseriforms also have 6–8 *MYH* genes, including *MYH13*, but not a functional or complete *MYH3*. Neoaves can have 8–12 *MYH* genes, usually including complete *MYH3* and *MYH13* coding sequences, but sometimes missing one or both. Passerines consistently have 10–12 *MYH* genes, due largely to a duplication of *MYH22* and *MYH23* that is conserved across the group. We label these copies *MYH22B/C* and *MYH23B/C*. They consistently appear in the order *MYH22B-MYH23B-MYH22C-MYH23C*, which likely indicates an unequal crossing over where both genes were duplicated together.

The subfamilies *MYH20* and *MYH21* are closely related in sequence and almost always single-copy. The phylogenetic relationships indicate one likely derived from duplication of the other. *MYH25* and *MYH26* at the opposite end of the cluster have the same relationship. The subfamilies *MYH22* and *MYH23* vary in number between 1–4 genes. *MYH24* is a small subfamily that does not appear in ratites or passerines and may be an offshoot of *MYH23*, that was lost in passerines. Some copies of *MYH22* and *MYH23* appear to be conserved duplicates among genera and families, while others appear to be recent species-specific duplications with >99% nucleotide sequence similarity.

**3.3) Crocodilia**

Crocodilians have 3–4 *MYH* genes in their *GAS7*-linked cluster in the order *MYH30-MYH31-MYH32-MYH33*, and do not have *MYH3* or *MYH13*. No species-specific additional duplicates were observed.

**3.4) Testudines**

Turtles and tortoises have 2–3 *MYH* genes in their *GAS7*-linked cluster as *MYH3-MYH35-MYH36-MYH37-MYH13*, with *MYH37* sometimes absent. No species-specific additional duplicates were observed.

**3.5) Squamata**

Snakes typically have the configuration *MYH3-MYH40-MYH41-MYH42* and lack *MYH13*. Lizards have 4–7 *MYH* core genes in the same order, but including *MYH13* and sometimes additional recent duplicates of *MYH40/MYH41/MYH42*.

**3.6) Anura and Caudata**

Anurans and Caudates have more complex multi-cluster structure compared to Amniote single core skeletal *MYH* configuration. Anurans and Caudates have *GAS7*-linked clusters that have *MYH3-MYH45* followed by 1–7 *MYH13*-subfamily genes, and ending (usually) with a single *MYH46*. Expression data indicates these are their primary skeletal muscle myosins and therefore the lone amniote MYH13 gene is either (1) a last remnant gene of a larger ancestral tetrapod set of *MYH13* genes that have been retained in amphibians or (2) amphibians have expanded MYH13. Anuran MYH13A and MYH13B proteins form a clade sister to caudate MYH13C, and the common MYH13A/B/C ancestor is related to the amniote MYH13 clade. We observe that MYH13A, MYH13B, and MYH13C show evidence of phylogenetic structure within species and turnover within each group that indicate a pattern of recent duplication-loss turnover, and therefore a determination between these two scenarios may not be possible. We have labeled the Anura and Caudata *MYH13* primary skeletal genes as *MYH13B* and *MYH13C*, respectively, to distinguish them from the specialized lone amniote *MYH13* gene.

MYH45 appears phylogenetically related to other tetrapod core MYH proteins in the expected phylogenetic position. Curiously, MYH46 is highly differentiated and phylogenetic appears to be related to Chondrichthyes MYH56 proteins. Therefore, MYH46 may be a much earlier derived protein that was never replaced in Amphibia, in the same way that *MYH13* has remained present in a single copy at the same distal 3'-end position of the cluster in Amniotes.

Both Anurans and Caudates have an additional tandem cluster of 1 (*Spea bombifrons*) to 15 (*Ambystoma mexicana*) MYH proteins labeled *MYH47A*. We refer to this cluster as the *TBC1D25*-linked cluster due to its consistent presence next to this gene. This cluster also appears as a single pseudogene in platypus (*Ornithorhynchus anatinus*) and a three-MYH cluster in West African lungfish (*Protopterus annectens*).

Anurans have an additional *TKTL2*-linked cluster of 2–4 genes labeled *MYH47B*, whose phylogenetic relationships indicate they are derived from MYH47 proteins. However, no one-to-one orthologs of *MYH47A* and *MYH48B* genes are indicated, indicating separate duplicative expansions or active turnover at both tandem cluster loci.

##### 3.7) Caecilia

Caecilians have one of the shortest *GAS7*-linked MYH clusters of any sarcopterygian group we observed, with a conserved configuration of only *MYH3-MYH45*. The lack of MYH13 genes in Caecilians, but their presence in anurans, caudates, and amniotes indicates a likely loss of *MYH13* subfamily genes in an ancestor of all extant caecilians. *Rhinatrema bivittatum* has a *TBC1D25*-linked cluster with two *MYH47A* genes, but *Microcaecilia* and *Geotrypetes* had no *MYH* genes at this locus. None of the caecilian genomes examined had a *TKTL2*-linked *MYH47B* cluster.

##### 3.8) Dipnoi

West African lungfish (*Protopterus annectens*) has a *GAS7*-linked cluster of 7 full-length and 2 pseudogenized *MYH* genes that are labeled *MYH50A1–MYH50A9*. They appear in linear order, with a possible *MYH3* gene at the proximal 5' end of the cluster next to *SCO1*. The current assembly of *Protopterus annectens* has a small gap where *MYH3* is located and while fragments can be seen on either side of the gap, the completeness or functionality of *MYH3* lungfish is indeterminate.

##### 3.9) Coelacanth

The West Indian Ocean coelacanth (*Latimeria chalumnae*) has a *GAS7*-linked cluster in the order *MYH3-MYH49A-MYH49B*. We note that the *MYH49* sequences are more closely related phylogenetically to tetrapods than lungfish *MYH48*, but after the divergence of *MYH45* in amphibians. This furthers the pattern that there seemed to be a mix of *MYH* ages in Sarcopterygian genomes, where we see a variety of phylogenetic relationships and divergence orders among *MYH45, MYH46, MYH48*, *MYH49*, and the *MYH13* subfamilies.

##### 3.10) Non-Teleost Actinopterygii

Ray-finned fish have an immense diversity of clusters, subfamilies, and paralog numbers that has been the subject of much of the previous work on MYH diversity (McGuigan et al. 2004, Ikeda et al. 2007). Substantial additional work will be needed beyond this to establish the history and diversity of MYH genes in this group. The complexity of the number of clusters and development of new subfamilies is closely tied to the key genomic transitions within this group. Grey bichir (*Polypterus senegalus*) has a single cluster at the *GAS7*-linked locus, with configuration *MYH3-MYH50A1-MYH50A2-MYH50A3-MYH53*. *MYH3* was apparently lost in most other species Actinopterygii we examined except for Clupeidae (herrings, sardines, and shads). This includes reedfish (*Erpetoichthys calabaricus*) and means MYH3 loss appears to have happened repeatedly in Actinopterygii. Spotted gar (*Lepisosteus oculatus*) similarly has a single *GAS7*-linked cluster with *MYH50A1-MYH51A1-MYH51A2-MYH52A1-MYH52A2-MYH52A3-MYH53-MYH53B 2*. The bowfin (*Amia calva*) shows a similar configuration, but apparently has expanded its GAS7-linked cluster of MYHs independently. egWe note that the *MYH53* subtype genes are consistently on the opposite strange in orientation relative to the other genes in cluster.or

Acipenseridae appear to have two *GAS7*-linked clusters due to whole genome duplication (WGD). For example, sterlet (*Acipenser ruthenus*) has *MYH52A1-MYH53* between *SCO1* and *GAS7* on Chromosome 17 and a single *MYH52B1* between *SCO1* and *GAS7* on Chromosome 19. Additionally, sterlet has a set of five tandem clusters of *MYH54* on the duplicated Chromosomes 57 and 58. These clusters show one-to-one relationships indicating duplications of the clusters after duplication of the paralogs into a tandem arrays.

**3.11) Teleostei**

Teleost fish are particularly diverse in their MYH configurations, and we only incompletely document them here. We denote the *MYH55* and *MYH56* subtypes as teleost-specific with lettered subtypes indicating different duplicated/translocated clusters.es*lettt* Torafugu (*Takifugu rubripes*) and yellowfin tuna (*Thunnus albacares*)*a* have a single *MYH51D* gene between *SCO1* and *GAS7*. Zebrafish (*Danio rerio*) has only a partial *MYH16* sequence between *gas7b* and *map24k4b* on Chromosome 12. *MAP24K4B* is near *SCO1* in syntenic order in most vertebrate genomes. *Takifugu* and *Danio* share two syntenic clusters of *MYH56B* and *MYH55C* that indicate these are conserved, but *Takifugu* has an additional cluster that is not present in *Danio*. Other species and groups also appear to have other tandem clusters in various genomic locations that are not necessarily syntenic with other groups. Notably, the three-spined stickleback (*Gasterosteus aculeatus aculeatus*) has a pair of *MYH55* genes on theit's Y chromosome, which is one of the only observed sex-chromosome-linked MYH genes. A fuller accounting of these clusters would be the subject of a research effort of its own. We note one other curiosity, which is the presence of a tandem cluster of 30 *MYH55* genes on Chromosome 14 in sardine (*Sardina pilchardus*), which was the most genes in a single cluster we observed. This appears to be specific to this species.

**3.12) Chondrichthyes**

Cartilaginous fish have a *GAS7*-linked cluster with 4–6 copies of what we label the *MYH57* subfamily. Elasmobranch sharks have *MYH3* at the proximal end of the cluster, but rays and the Australian ghost shark (*Callorhinchus milii*) lack *MYH3*.

**3.13) Lampreys**

Sea lamprey (*Petromyzon marinus*) and Far Eastern brook lamprey (*Lethenteron reissneri*) each have a *GAS7*-linked cluster including three genes from the *MYH58* subfamily. We note that *SCO1* does not appear at the 5′ end of the cluster, but instead *MAP24K4B* which is commonly in the gene order further 5′ in most other vertebrates and is also the proximal gene to the cluster in some Actinopterygii.

**3.14) Hagfish**

The Atlantic hagfish (*Myxine glutinosa*) has a cluster including two genes from the *MYH57* subfamily. The neighboring genes are *SDHD* and *KIAA0513* to this cluster, which may indicate a translocation in the hagfish.

**3.15) Lancelets**

Lancelets from *Branchiostoma* have two tandem *MYH* clusters in different genomic locations, containing 5–9 *MYH* genes each. Neither cluster appears to be linked to *GAS7* nor any other gene from that neighborhood. The sequences of the MYH proteins are highly divergent even compared to tunicates and cause issues with both alignment and phylogenetic inference of vertebrate sequences. This is not necessarily unexpected for lancelets, which have experienced a long evolutionary timeline apart from other vertebrates and have a divergent physiology. We have set these sequences aside for further study later, and for the present we document here the presence of tandem *MYH* clusters in this lineage.

**3.16) Tunicates**

The sea squirt *Styela clava* and vase tunicate *Ciona intestinalis* have a tandem cluster of 2–6 *MYH59A and MYH59B* genes (with many apparent pseudogenes) and a single *MYH59C* at a separate locus. These clusters are not linked to *GAS7* or any other syntenic locus with vertebrates. These two gene families do not share the closest common ancestor with the vertebrate core skeletal muscle myosins, nor are they close in sequence with lancelets. Instead, these two types appear to diverge before the divergence of *MYH7B* and *MYH15* lineage.

**4) Additional notes on gene conversion and intergenic recombination.**

The coding sequences of monotremes, marsupials, and placental mammals are known to have markedly different biases in the GC content of the third codon position or $GC_3$ content [16,17]. Since MYH protein sequences are highly conserved generally even across the mammalian myosin forms, the substantial shift in $GC_3$ content bias was sufficient in the case of monotremes to make their MYH protein sequences appear monophyletic within monotremes. The $GC_3$ content bias is easily visually apparent (Fig. S3). However, when we model the phylogeny removing third positions, we can see each ortholog of *MYH1, MYH2, MYH4,* and *MYH8* return to their expected phylogenetic position. Pairwise distance analysis of the protein sequences between any combination of platypus and echidna with human and elephant shows that genes have one-to-one similarity in the same linear order in the cluster. We conclude that all mammals including monotremes have an orthologous set of MYH proteins and that the apparent monophyly of monotreme sequences is caused by the dramatic $GC_3$ content shift. We did not observe this phenomenon in any other vertebrate group that would indicate composition shifts are creating the non-orthologous phylogenetic patterns between major vertebrate groups. In summary, monotreme sequences at first appeared to have also undergone a loss-duplication shift from other mammals, but this was due only to $GC_3$ content shifts, not gene replacement.

Several instances of recombinations among neighboring MYH genes were apparent in mammals and birds (Fig. S4). The alignments clearly show regions that are partial gene recombination events likely also due to unequal crossing over that occurred inside an MYH locus rather than between copies of MYH. In general, this seemed to be a more minor contributor than whole gene replacement, based on the evidence available. A more comprehensive examination of this phenomenon will be necessary to determine the rate. We do not find that this process alone appears to be the primary driver for the lack of orthology between groups. The much more common presence of recent whole-gene duplicates and full-length pseudogenes indicates this is a less frequent contributor.

We also note that gene conversion, GC-bias, and partial recombinant genes will all be overwritten by whole gene loss and duplications, so we both expect and observe that these phenomena are more rare than whole gene unequal crossing over as a contributor to our overall model of MYH evolution.

**5) Additional notes on extreme muscle MYH expression**

Extreme and specialized muscles with unusually fast rates of contraction also bear some reconsideration in light of this model and our findings. We analyzed several muscles that are considered specialized and extreme, including rattlesnake caudal muscles, hummingbird pectoral, manakin scapulohumeralis, and syringeal muscles in a finch. In each case, different MYH proteins were predominantly expressed, meaning there is not a universal extreme muscle myosin but instead multiple evolutions of MYH molecules that might contribute to high contractile speeds. MYH13 is commonly expressed in mammalian laryngeal and extraocular tissues [18]. Our data and previous studies support that MYH13 is a key sarcomeric myosin in avian syringeal muscles, though it does not appear to be the sole MYH expressed [19]. These muscles typically generate superfast contractions with frequencies greater than 90 Hz [20]. It has been suggested that MYH13 is limited in that it contributes to high contractile speeds but only in relatively small muscles with little force requirements [18]. MYH26 was also expressed at high levels in *Lonchura* syrinx tissue, and its ortholog in hummingbird was expressed in the tongue but not in locomotor muscles. While MYH13 is undoubtedly expressed in high amounts in muscles used in vocalization, more research is needed to understand how MYH26 may also contribute to muscles with specialized contractile needs in birds.

**6) Additional discussion on the potential impact of MYH diversity relative to other muscle factors**

How much does MYH diversity determine muscle performance compared to other factors? MYH proteins are ultimately the molecule that provides the motor force by motion of the myosin heads, but certainly other molecules might provide additional performance modifications. We can first consider other fibers, specifically actin and titin. Titin is a single-copy gene in all sarcopterygian genomes where it is present, though is present in higher copy numbers in teleosts due to whole-genome duplication events. Sarcomeric actin is present in all the vertebrate genomes we analyzed in exactly three copies: skeletal muscle *ACTA1*, smooth muscle *ACTA2*, and cardiac *ACTC1*. Our expression datasets also showed only expression of *ACTA1* in skeletal muscles uniformly. Sequence variants in actin and titin could be impacting performance, but since they are single-copy, these changes would impact all muscles. Therefore, even if actin or titin is contributing to the interspecific or intergroup differences in muscle performance, it cannot account for the various muscle performance properties within a species.

Myosin light chain (MYL) proteins are another molecule closely tied to contractile function and the myosin head. We did identify MYL binding sites on MYH proteins as a potential hotspot of repeated diversification. In general, skeletal muscle-associated MYLs are fewer and less variable in paralog numbers than skeletal muscle MYHs. MYLs also appear individually and do not occur in a variable tandem cluster. MYLs remain an interesting candidate for further study.

The other system that is doubtlessly a major player in modulating muscle performance properties is the calcium cycling system, including the SERCA pump and ryanodine receptor release channels for $Ca^{2+}$ ions. This system has undergone multiple evolutionary transitions both in the form of specific molecules like calsequestrin [21] or the overall cell and tissue architecture. While the calcium-cycling system sets the rate at which excitation-contraction cycle can happen, the synchronicity and motor force of the muscle contractions themselves is considered more driven by the structure and action of the fiber proteins themselves. Again, as with the other fiber molecules discussed, these proteins tend to be single or few in copy number and the major players in this system have not been necessarily shown to vary substantially among locomotor skeletal muscles.

We conclude that MYH genes are more variable in copy number and diverse in function than most other key parts of this system. The evidence seems to point toward MYH molecular diversity and use ratios being a potentially sizable portion of the functional variation influencing the muscle performance phenotypes. Additionally, work will be needed to test these hypotheses, as we further understand the biomechanical, physiological, cellular, and molecular layers of the evolution of vertebrate skeletal muscles.

### Supplementary Figures

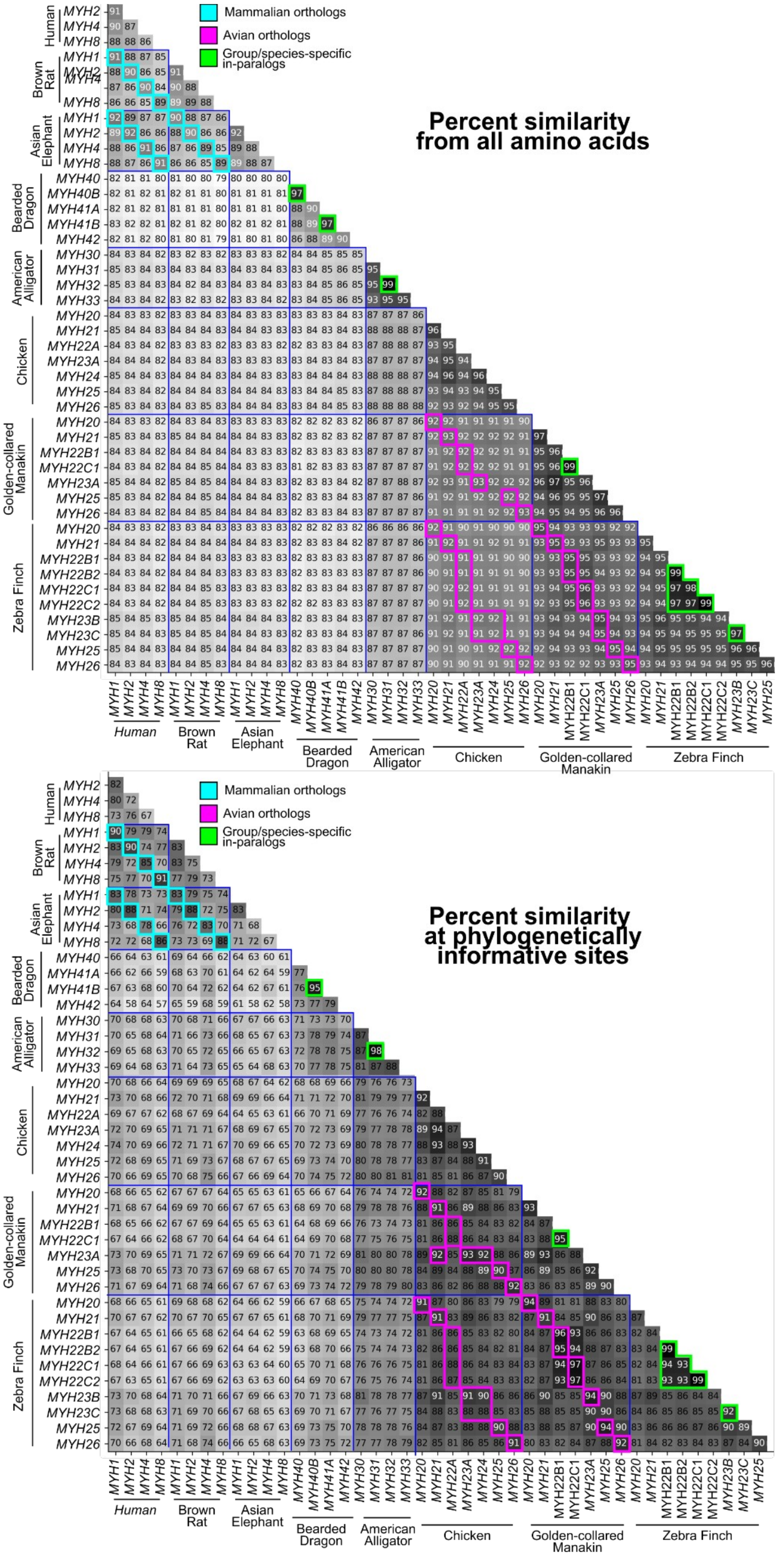


**Figure S1.** Pairwise sequence distances from selected amniotes show the presence of intragroup, but not intergroup orthologs.

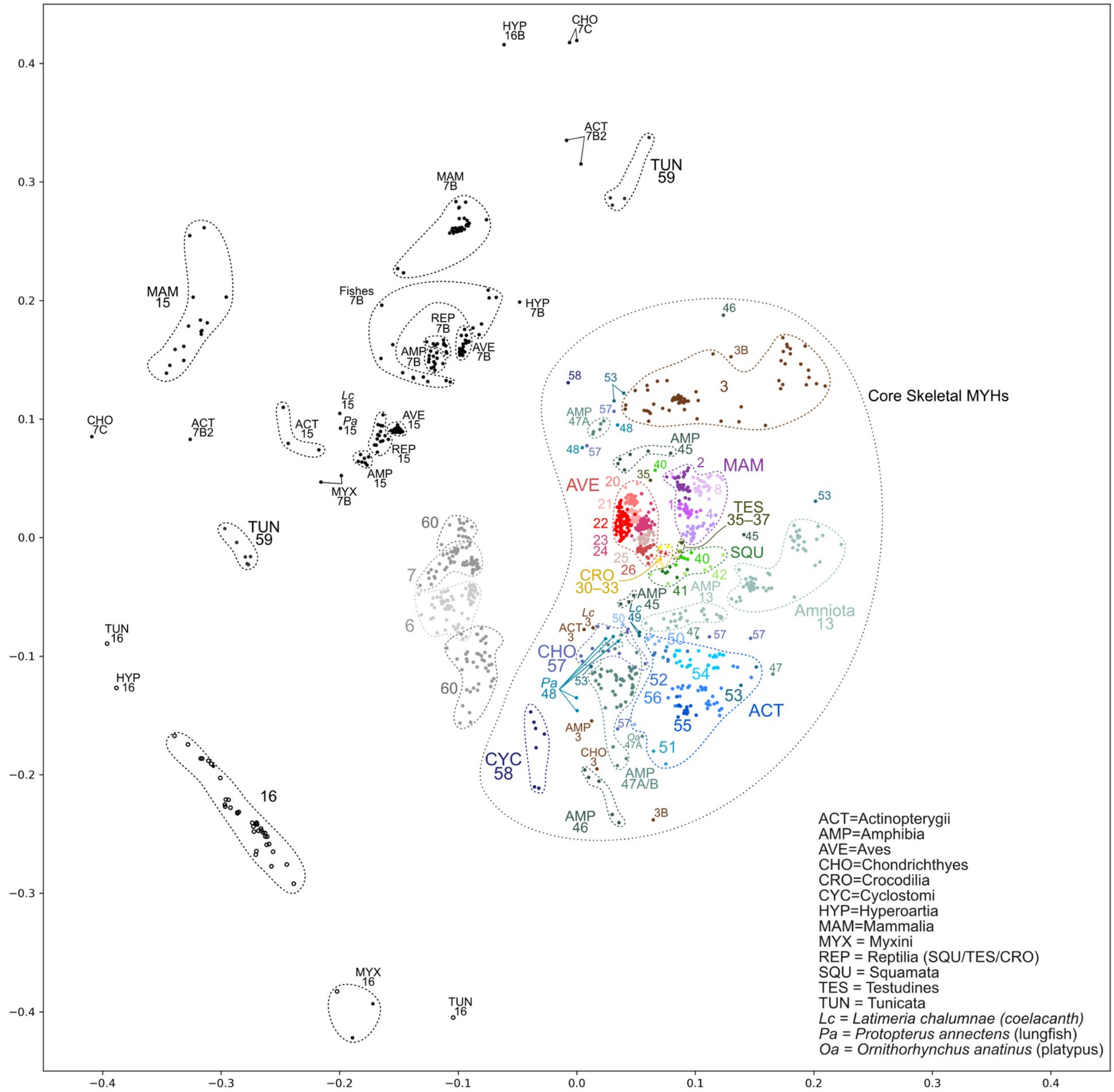


**Figure S2.** Multidimensional scaling plot from pairwise distances of amino acid sequences for all *MYH* genes in the primary alignment. Groups and outlier genes are labeled with their MYH number, according to our revised naming scheme.

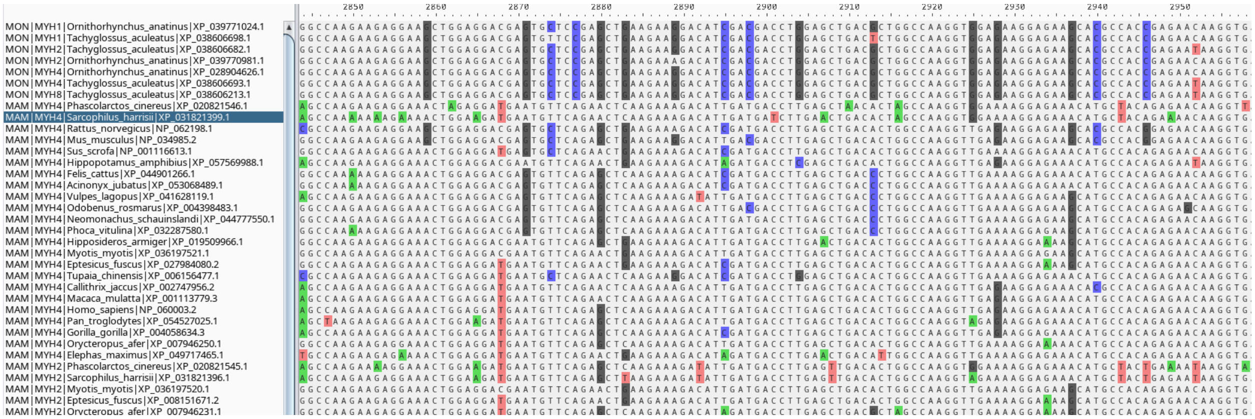

**Figure S3.** Example alignment of mammalian *MYH* codon sequences showing the GC3 bias in monotremes.

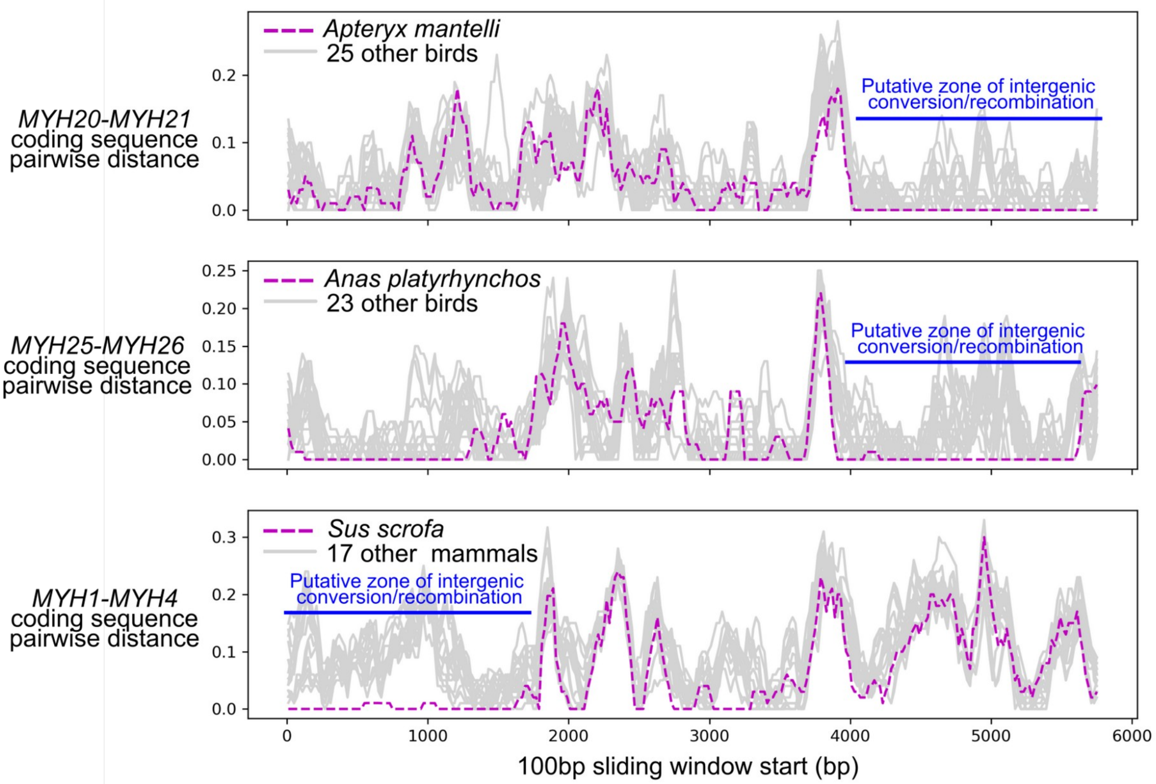


**Figure S4.** Three examples of potential intergenic recombination or gene conversion based on 100bp sliding-window nucleotide pairwise distance from the coding sequences of adjacent gene pairs. Examples show: apparent replacement genomic sequence in *Apteryx MYH21* tail region in with identical sequence from *MYH20* (top), the *Anas MYH26* tail region from *MYH25* (middle), and the *Sus MYH4* head region from *MYH1*.

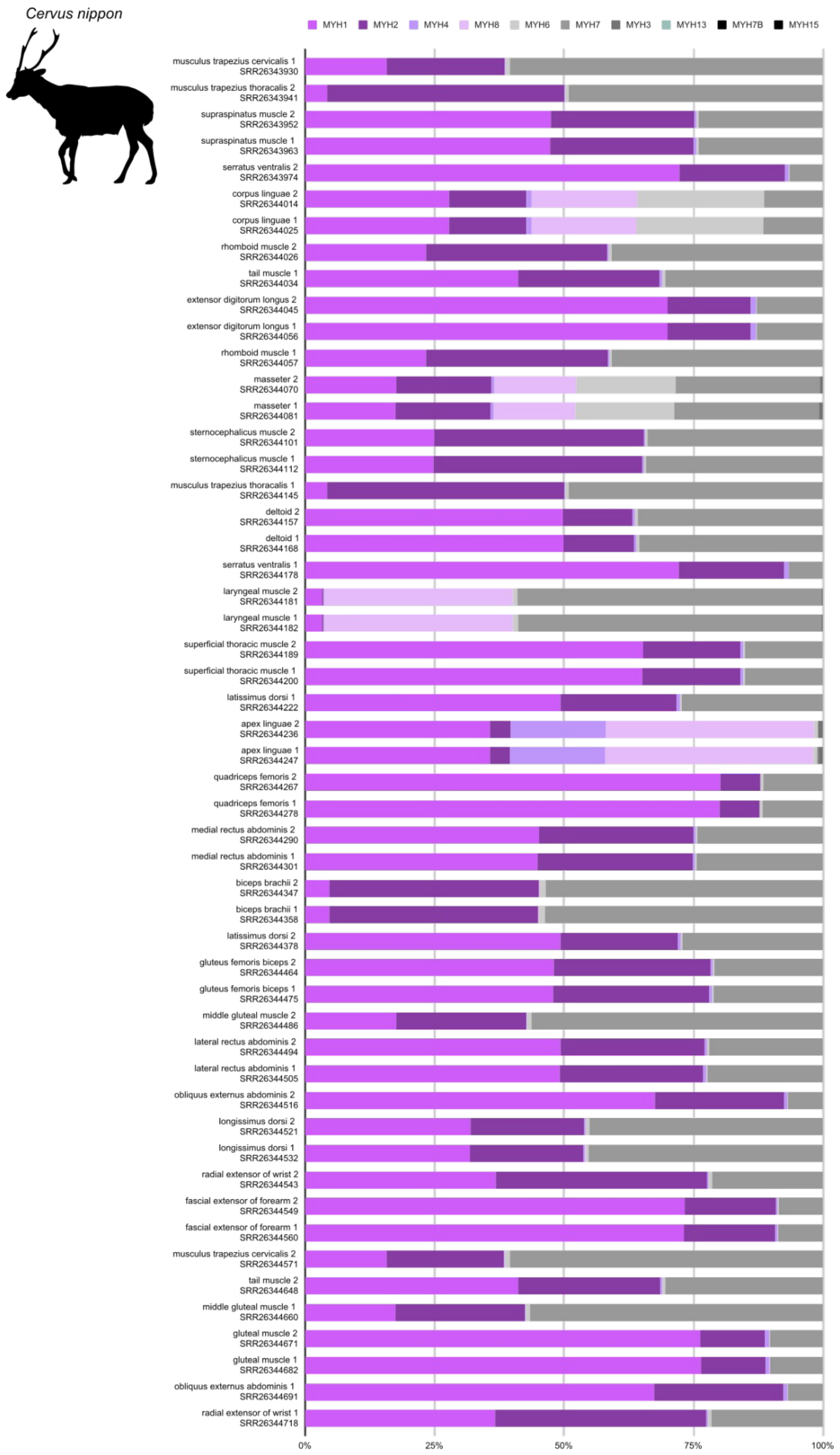


**Figure S5.** Sarcomeric MYH expression relative proportions in 24 sika muscle tissues from two individuals.

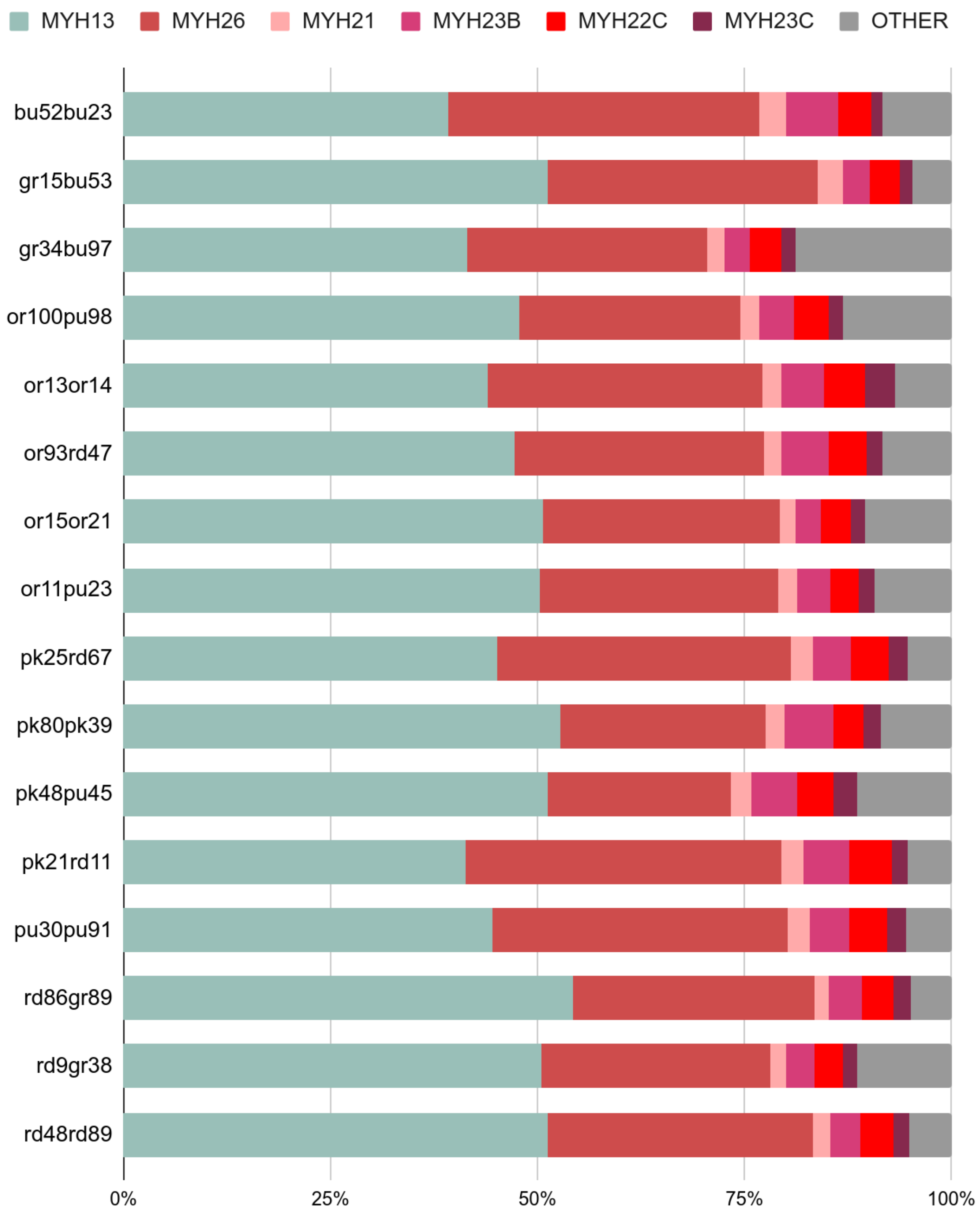


**Figure S6. MYH expression relative proportions in Lonchura striata syrinx muscle tissues from sixteen individuals.**

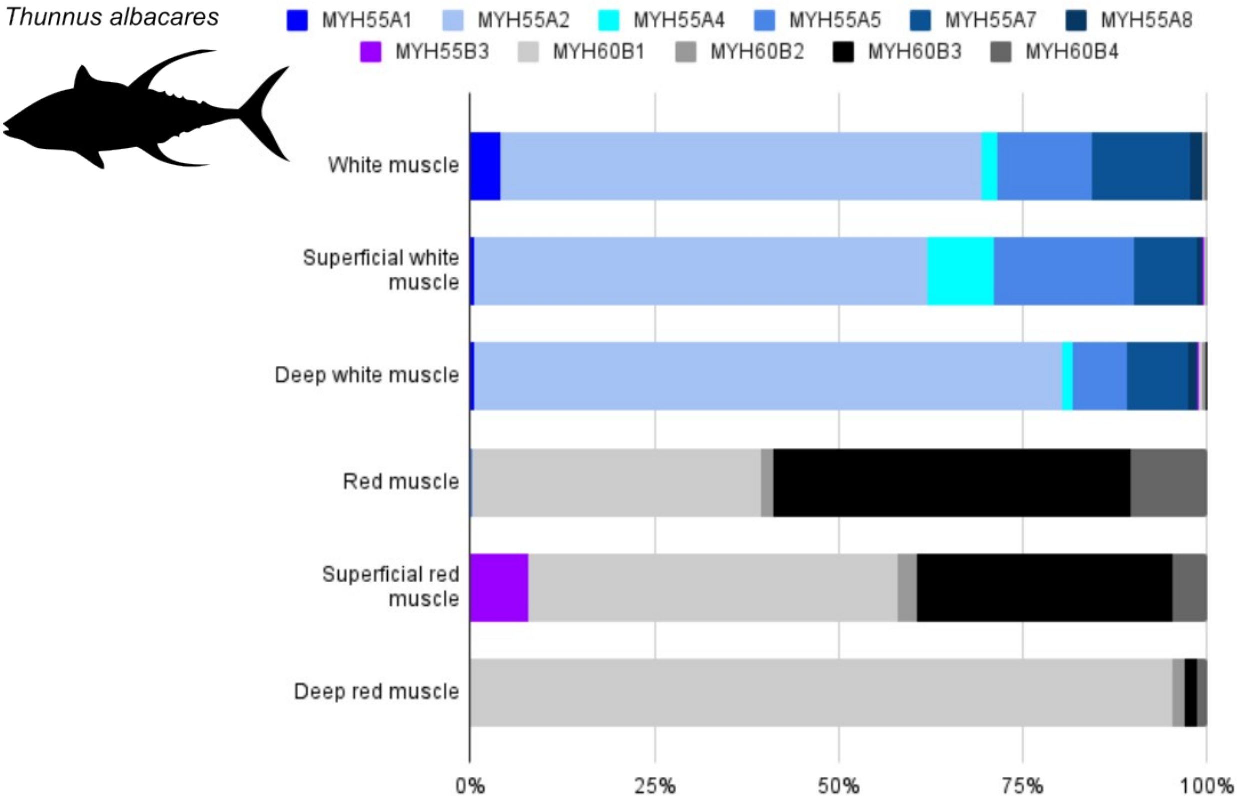


**Figure S7.** Sarcomeric MYH expression relative proportions in 6 muscle tissues from individuals (1 and 2).

### Supplementary Data

**Data S1. FASTA file containing complete sequence alignment used to infer pairwise distances and loop lengths.**

**Data S2. FASTA file containing trimmed alignment used for phylogeny.**

**Data S3. Phylogenetic tree in Newick format with bootstrap support values.**

**Data S4. Spreadsheet containing description of sequences included in Data S1. Also included are comments on removed or edited sequences.**

**Data S5. Spreadsheet containing accession numbers to data sets and raw count values used to calculate expression for Figures 1 and 3.**

**Data S6. FASTA file containing sequence alignment of tetrapod core cluster myosins used to generate conservation scores.**

**Data S7. CSV containing calculated conservation scores of tetrapod core cluster MYHs.**

**Data S8. FASTA file containing coding sequences for sliding window analysis in Figure S4.**

#### Supplementary References

1. Tajsharghi H *et al.* 2014 Recessive myosin myopathy with external ophthalmoplegia associated with MYH2 mutations. *European Journal of Human Genetics* **22**, 801–808. (doi:https://doi.org/10.1038/ejhg.2013.250)

2. Scala M, Accogli A, Grandis ED, Allegri A, Bagowski CP, Shoukier M, Maghnie M, Capra V. 2018 A novel pathogenic *MYH3* mutation in a child with Sheldon–Hall syndrome and vertebral fusions. *American Journal of Medical Genetics Part A* **176**, 663–667. (doi:https://doi.org/10.1002/ajmg.a.38593)

3. Finno CJ *et al.* 2018 A missense mutation in MYH1 is associated with susceptibility to immune-mediated myositis in Quarter Horses. *Skeletal Muscle* **8**, 7. (doi:https://doi.org/10.1186/s13395-018-0155-0)

4. Johnson CA *et al.* 2021 Identification of sequence changes in myosin II that adjust muscle contraction velocity. *PLOS Biology* **19**, e3001248. (doi:https://doi.org/10.1371/journal.pbio.3001248)

5. Lindqvist J, Iwamoto H, Blanco G, Ochala J. 2013 The fraction of strongly bound cross-bridges is increased in mice that carry the myopathy-linked myosin heavy chain mutation MYH4L342Q. *Disease Models & Mechanisms* **6**. (doi:https://doi.org/10.1242/dmm.011155)

6. Uctepe E, Mancılar H, Esen FN, Unverengil GG, Vona B, Yesilyurt A. 2025 A homozygous myh1 variant underlies autosomal recessive isolated recurrent rhabdomyolysis. *American journal of medical genetics. Part A* **197**, e63952. (doi:https://doi.org/10.1002/ajmg.a.63952)

7. Veugelers M *et al.* 2004 Mutation of perinatal myosin heavy chain associated with a carney complex variant. *The New England Journal of Medicine* **351**, 460–469. (doi:https://doi.org/10.1056/nejmoa040584)

8. Friedman CE *et al.* 2024 Multiplexed functional assessments of MYH7 variants in human cardiomyocytes. *Circulation: Genomic and Precision Medicine* **17**. (doi:https://doi.org/10.1161/circgen.123.004377)

9. Tajsharghi H, Darin N, Rekabdar E, Kyllerman M, Wahlström J, Martinsson T, Oldfors A. 2005 Mutations and sequence variation in the human myosin heavy chain IIa gene (MYH2). *European Journal of Human Genetics* **13**, 617–622. (doi:https://doi.org/10.1038/sj.ejhg.5201375)

10. Meredith C *et al.* 2004 Mutations in the slow skeletal muscle fiber myosin heavy chain gene (MYH7) cause Laing early-onset distal myopathy (MPD1). *The American Journal of Human Genetics* **75**, 703–708. (doi:https://doi.org/10.1086/424760)

11. Laing NG *et al.* 1995 Autosomal dominant distal myopathy: linkage to chromosome 14. *The American Journal of Human Genetics* **56**, 422–427.

12. Hedera P, Petty EM, Bui MR, Blaivas M, Fink JK. 2003 The second kindred with autosomal dominant distal myopathy linked to chromosome 14q. *Archives of Neurology* **60**.

(doi:https://doi.org/10.1001/archneur.60.9.1321)

13. Dye DE, Azzarelli B, Goebel HH, Laing NG. 2006 Novel slow-skeletal myosin (MYH7) mutation in the original myosin storage myopathy kindred. *Neuromuscular Disorders* **16**, 357–360. (doi:https://doi.org/10.1016/j.nmd.2006.03.011)

14. Ikeda D, Ono Y, Snell P, Edwards YJK, Elgar G, Watabe S. 2007 Divergent evolution of the myosin heavy chain gene family in fish and tetrapods: evidence from comparative genomic analysis. *Physiological Genomics* **32**, 1–15. (doi:https://doi.org/10.1152/physiolgenomics.00278.2006)

15. Schiaffino S, Hughes SM, Murgia M, Reggiani C. 2023 MYH13, a superfast myosin expressed in extraocular, laryngeal and syringeal muscles. *The Journal of Physiology* **602**, 427–443. (doi:https://doi.org/10.1113/jp285714)

16. van Rheede T, Bastiaans T, Boone DN, Hedges SB, de Jong WW, Madsen O. 2005 The platypus is in its place: nuclear genes and indels confirm the sister group relation of monotremes and therians. *Molecular Biology and Evolution* **23**, 587–597. (doi:https://doi.org/10.1093/molbev/msj064)

17. Romiguier J, Ranwez V, Douzery EJP, Galtier N. 2010 Contrasting GC-content dynamics across 33 mammalian genomes: Relationship with life-history traits and chromosome sizes. *Genome Research* **20**, 1001-1009. (doi:https://doi.org/10.1101/gr.104372.109)

18. Schiaffino S, Hughes SM, Murgia M, Reggiani C. 2024 MYH13, a superfast myosin expressed in extraocular, laryngeal and syringeal muscles. *The Journal of Physiology* **602**, 427-443. (doi:https://doi.org/10.1113/JP285714)

19. Mead AF et al. 2017 Fundamental constraints in synchronous muscle limit superfast motor control in vertebrates. eLife 6. (doi:https://doi.org/10.7554/elife.29425)

20. Rome L. 2006 Design and function of superfast muscles: new insights into the physiology of skeletal muscle. *Annual Review of Physiology* **68**, 193-221 (doi:https://doi.org/10.1146/annurev.physiol.68.040104.105418)

21. Harvey CM, Fuxjager MJ, Pease JB. 2025 Deep-time gene expression shift reveals an ancient change in avian muscle phenotypes. *PLoS Genetics* **21**, e1011663–e1011663. (doi:https://doi.org/10.1371/journal.pgen.1011663)
